# IL12-engineered human PSMA-CAR T cells for the treatment of advanced prostate cancer

**DOI:** 10.64898/2026.03.05.709907

**Authors:** Lupita S. Lopez, Ziyou Cui, Yukiko Yamaguchi, John P. Murad, Zhiyuan Yang, Ke Zou, Jason Yang, Wen-Chung Chang, Stephen J. Forman, Vivien W. Chan, Saul J. Priceman

**Affiliations:** Division of Medical Oncology, Department of Medicine, Keck School of Medicine (KSOM) of USC, Los Angeles, CA, 90033, USA; Irell and Manella Graduate School of Biological Sciences, Beckman Research Institute of City of Hope, Duarte, CA, 91010, USA; Department of Hematology and Hematopoietic Cell Transplantation, City of Hope, Duarte, CA, 91010, USA; Eureka Therapeutics, Inc., Emeryville, CA, 94608, USA; KSOM/Norris Center for Cancer Cellular Immunotherapy Research, USC, Los Angeles, CA, 90033, USA

**Keywords:** chimeric antigen receptor, prostate cancer, adoptive cellular immunotherapy, prostate specific membrane antigen

## Abstract

Adoptive cell therapies used to treat advanced prostate cancer are being developed to target several tumor-associated antigens, including prostate-specific membrane antigen (PSMA). Chimeric antigen receptor (CAR) T cell therapy using the single chain variable fragment (scFv) derived from the humanized murine mAb clone, J591, as the antigen-binding domain has shown promising anti-tumor activity. However, it has also been associated with macrophage activation syndrome and other unwanted toxicities, highlighting the need for more specific and human-derived antigen-binders with optimized construct designs for improved safety and efficacy. Here, we optimize a human scFv-based PSMA-targeted CAR (hPSMA-CAR) with highly selective PSMA targeting. We further introduce a membrane-bound IL-12 (mbIL12) molecule, which enhances potency with increased T cell expansion, IFNy production and anti-tumor cell activity *in vitro*. Using two clinically-relevant bone-metastatic prostate cancer models, we show that mbIL12-engineered hPSMA-CAR T cells drive potent *in vivo* anti-tumor responses. In summary, we have developed a promising therapeutic that has potential to promote safe and effective treatment of advanced PSMA+ prostate cancer.

## Introduction

Metastatic castration-resistant prostate cancer (mCRPC) represents a challenging stage of prostate cancer with a 5-year survival rate around 30% and limited effective treatment options^1^. Although immune checkpoint blockade (ICB) has had striking clinical successes in select solid and hematological malignancies, these therapies have not shown similar therapeutic responses in mCRPC^2^. Therefore, bispecific T cell engagers (BITEs) and chimeric antigen receptor (CAR) T cell therapies have been an active area of clinical investigation^3–7^. Several tumor targets are attractive in mCRPC, including prostate specific membrane antigen (PSMA)^8^. Several groups have reported on clinical experiences with CAR T cells targeting PSMA, which have to date been challenged with severe adverse events^6,9,10^. As such, there is a critical need for potent, and safe, PSMA-targeted CAR T cell therapies.

Here, we report on the systematic optimization of a human PSMA (hPSMA) CAR T cell for the treatment of mCRPC. Using Eureka Therapeutics’ E-ALPHA^®^ phage display library, we identified a lead human anti-PSMA scFv, optimized the CAR extracellular spacer and transmembrane domains for selective PSMA targeting. We found that a dCH2 spacer together with a CD28 transmembrane domain on a 4-1BBz-containing CAR showed the greatest specificity for PSMA-expressing tumor cells. To address suboptimal potency of the hPSMA-CARs, scFv affinity maturation was performed, but did not significantly improve potency of hPSMA-CAR T cells. We then evaluated engineering hPSMA-CAR T cells with a membrane-bound IL12 (mbIL12) molecule to enhance anti-tumor activity. hPSMA-CAR/mbIL12 T cells showed improved tumor cell killing, expansion, and cytokine secretion, while maintaining highly specific PSMA targeting. Using an *in vitro* macrophage:tumor:T cell co-culture model, we show an improved cytokine safety profile of hPSMA-CAR/mbIL12 T cells compared with J591-based CAR T cells. Finally, we showed our hPSMA-CAR/mbIL12 T cells resulted in robust anti-tumor activity using *in vivo* and bone metastatic prostate cancer models. These data highlight PSMA-CAR T cells engineered with mbIL12 as a promising therapy for the treatment of mCRPC, with broader implications for the development of safe and effective CAR T cells for other solid tumors.

## Results

### Development of a human-derived PSMA-CAR T cell

We first attempted to develop a PSMA-CAR using a human scFv to reduce toxicity, and T cell clearance due to anti-scFv immunogenicity. Eureka Therapeutics’ E-ALPHA**^®^** Phage Display platform was used to screen anti-PSMA antibodies in single-chain variable fragment (scFv) format using PSMA over-expressing Jurkat cells. After three rounds of panning, 1080 clones were screened, and of these, two unique clones were obtained, ET260-1 and ET260-2. These clones were confirmed by DNA sequencing. Both clones stained PSMA positive cells but not PSMA negative cells (**Figure S1**). Binding kinetic studies of the ET260 clones showed ET260-2 had lower binding (on-rate) and dissociation (off-rate) compared to ET260-1 (**Figure S1, Table S1).**

The ET260-1 and ET260-2 scFvs were cloned into a CAR construct comprising a CH3-IgG extracellular spacer (CH2-deleted, termed dCH2), a CD4 transmembrane domain (CD4tm), an intracellular 4-1BB costimulatory domain (41BB), and a CD3z cytolytic domain (CD3z), along with a truncated CD19 gene (CD19t) separated by a T2A skip sequence (referred to as hPSMA-CARs) **(Figure 1a)**. The J591 scFv was cloned into the same construct to serve as a reference of activity for our hPSMA-CARs. T cells were transduced with lentivirus encoding the CAR constructs and enriched for CD19t (**Figure S2a**). We detected the CAR directly through the dCH2 extracellular spacer by anti-Fc staining and expression was comparable across the three PSMA-CARs (**Figure 1b**). We then tested these CARs against PC-3 cells in a coculture over 72 hrs at an effector:tumor cell (E:T) ratio of 1:4. These cells were engineered to express varying levels of PSMA (**Figure S3**). The ET260-1 CAR had greater tumor cell killing, activation, and IFNy secretion compared to the ET260-2 CAR (**Figure 1c-e**). While J591-based CAR T cells showed superior tumor cell killing, activation, and IFNy release against PSMA-expressing PC-3 cells compared with the hPSMA-CAR T cells, J591-CARs also had significantly higher tumor cell killing and activation against the PSMA-negative PC-3 parental cell line (WT) compared to the hPSMA-CARs (**Figure 1c,d**). For the purpose of our studies, we use activity against PC-3 WT as a proxy for on-target off-tumor toxicities due to the low levels of PSMA^11–14^ (**Figure S3**).

**Figure 1.**
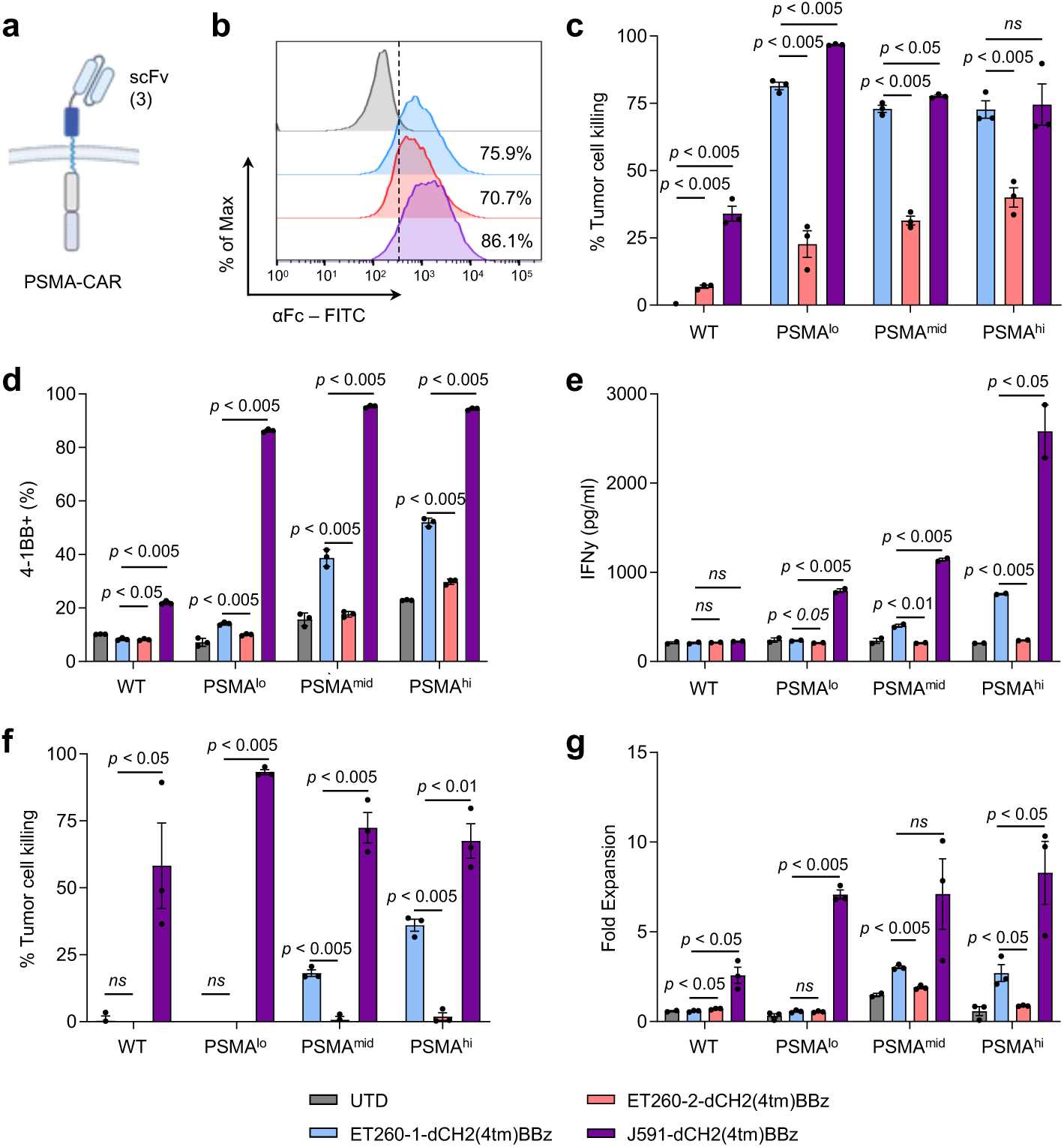
Development of a human PSMA-CAR T cell **a** Schema of hPSMA-CAR scFv variants with a dCH2 Fc extracellular linker, CD4tm, 4-1BB costimulatory domain, and CD3ζ signaling domain construct. **b** Flow cytometric analysis of CARs on the surface of primary T cells via detection of the Fc linker domain. **c** Tumor cell killing of hPSMA-CARs over 72 hrs when co-cultured with PSMA-expressing cell lines at an effector to tumor ratio (E:T) of 1:4. **d** Expression of 4-1BB on hPSMA-CARs compared to the J591-CAR after 72 hrs. **e** Production of IFNγ at 72 hrs as determined by ELISA. **f,g** Tumor cell killing and fold expansion of hPSMA CARs after 8-day LTK at E:T of 1:20. Data are presented as mean values ± SEM. *P* values indicate differences between ET260-1-dCH2(4tm)BBz and ET260-2-dCH2(4tm)BBz and between ET260-1-dCH2(4tm)BBz and J591-dCH2(4tm)BBz using a two-tailed Student’s t test.

To further assess functionality of these CARs, we performed extended long-term killing assays (LTK) over 8 days at an E:T of 1:20. The hPSMA-CARs did not perform equivalently to J591-CARs with lower tumor cell killing and fold expansion (**Figure 1f,g**). Importantly, the ET260-1 CAR showed promising effector function against PC-3 PSMA cell lines and limited activity against PC-3 WT and based on these results, the ET260-1 scFv was selected for further optimization. These studies suggest that while human scFv-based PSMA-CARs can target PSMA+ tumor cells *in vitro*, their activity was diminished compared with humanized J591-CAR. This may be explained by the binding kinetics of the J591, which showed higher binding (on-rate) and lower dissociation (off-rate) compared to the ET260 scFvs (**Table S1**).

### Optimization of hPSMA-CAR extracellular spacer and transmembrane domains

We next evaluated whether optimization of the CAR backbone with the ET260-1 scFv could improve the potency of hPSMA-CARs. We evaluated hPSMA-CAR constructs using three different extracellular spacer lengths (dCH2, CD8h, and HL) and three different transmembrane domains (CD4tm, CD8tm, and CD28tm), all using the BBz backbone (**Figure 2a**). T cells showed comparable transduction efficiency and were subsequently enriched for CD19t for further studies (**Figure S2b**). We observed higher Fc staining in the dCH2(28tm)BBz containing hPSMA-CAR compared to the original CD4tm construct and the J591-CAR (**Figure 2b**). To evaluate *in vitro* functionality of these CAR T cells, we performed tumor cell killing assays against PC-3 cell engineered with varying levels of PSMA for 72 hours. ET260-1-dCH2(4tm)BBz and ET260-1-dCH2(28tm)BBz CAR T cells performed comparably to J591-CAR T cells against PC3 PSMA^mid^ and PSMA^hi^ (**Figure 2c**). The dCH2(28tm) containing hPSMA-CAR T cells showed higher 4-1BB expression and IFNy secretion compared to the dCH2(4tm), CD8h(8tm) and HL(28tm) containing hPSMA-CAR T cells. These versions of hPSMA-CAR T cells showed minimal increases in non-specific activity against PC-3 WT cells, as compared to J591-CAR T cells (**Figure 2c,d**). hPSMA-CAR T cells containing dCH2(CD28tm) showed improved activity against our PC-3 PSMA cell compared to dCH2(4tm), but were still suboptimal compared to the J591-CAR T cells (**Figure 2f,g**). Although we saw suboptimal activity of our ET260-1-based CAR T cells compared to J591-CAR T cells, it is important to highlight the specificity and sensitivity to PSMA combined with limited activity to the WT cell line. In summary, we improved potency of the hPSMA-CAR T cells with dCH2(28tm)BBz, but was still suboptimal in long-term tumor cell killing potential compared with J591-CAR T cells.

**Figure 2.**
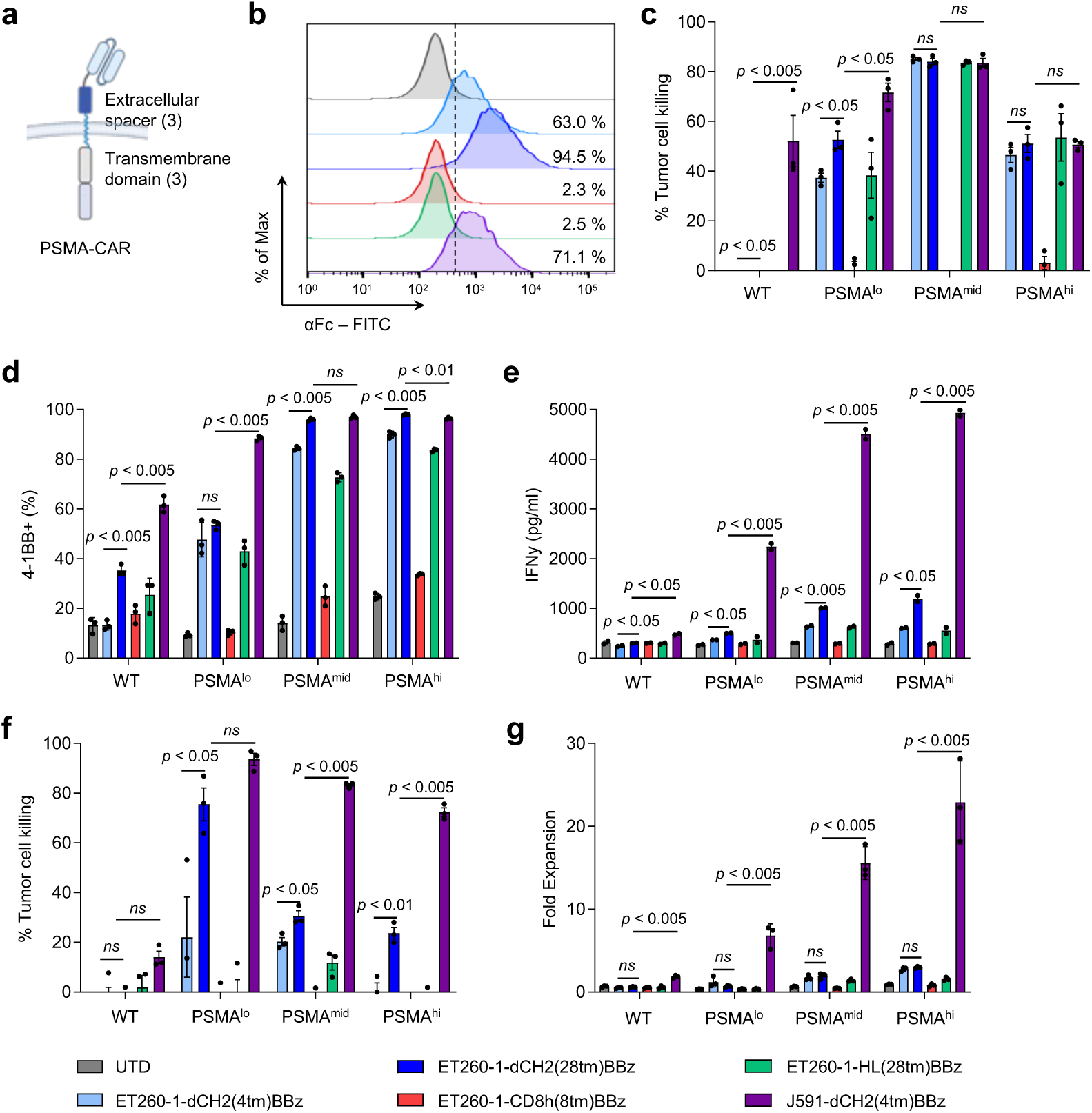
Optimization of hPSMA-CAR extracellular spacer and transmembrane domains **a** Schema of hPSMA-CAR variants with different extracellular linker (dCH2, CD8h, HL) and transmembrane domains (CD4tm, CD28tm, CD8tm), 4-1BB costimulatory domain, and CD3ζ signaling domain. **b** Flow cytometric analysis of ET260-1 CAR variants on the surface of primary T-cells via detection of Fc. **c** Tumor cell killing of hPSMA-CARs over 72 hrs when co-cultured with PSMA-expressing cell lines at an E:T of 1:4. **d** Expression of 4-1BB on ET260-1 CAR variants compared to J591 after 72 hrs. **e** Production of IFNy at 72 hr as determined by ELISA. **f,g** Tumor cell killing and fold expansion of ET260-1 CAR variants after 8-day LTK at E:T of 1:20. Data are presented as mean values ± SEM. *P* values indicate differences between ET260-1-dCH2(4tm)BBz and ET260-1-dCH2(28tm)BBz and between ET260-1-dCH2(28tm)BBz and J591-dCH2(4tm)BBz using a two-tailed Student’s t-test

### Affinity maturation of hPSMA scFv impacts CAR stability and activity in vitro

We hypothesized that the potent activity of J591-CAR T cells was in part due to its high affinity to PSMA. We therefore sought to improve the binding kinetics of our ET260-1 scFv through affinity maturation (AM). Indeed, we saw that while J591 had a high on-rate, it also had a particularly slow off-rate (**Table S1**). In contrast, our ET260-1 scFv had a lower on-rate and higher off-rate than J591. To address this, AM using error-prone PCR was performed, identifying 10 unique affinity-matured scFv clones carrying different mutations which were shown to bind Jurkat-PSMA cells but not Jurkat parental cells. These clones were subjected to Surface Plasmon Resonance to determine affinity for the PSMA extracellular domain protein (**Figure S4**). The parental scFv, ET260-1(WT), served as a positive control (**Table 1**). In addition to ET260-1, J591 served as a reference of kinetic activity and based on the binding kinetics of the clones relative to both ET260-1 and J591, five clones were selected and cloned into our dCH2(CD28tm)BBz CAR construct. Specifically, clones were selected based on their improved KD compared to the parental ET260 and kinetics that resembled J591, i.e. higher on-rate and slow off-rate (**Table S3**). T cells were engineered to express CARs containing the respective scFv and enriched for CD19t. Interestingly, of the five clones, clones 56 and 69 did not have stable CAR expression (by Fc staining) while clone 39 had two distinct populations. Clones 58 and 67 most closely resembled the parental CAR expression pattern (**Figure 3a**).

**Figure 3.**
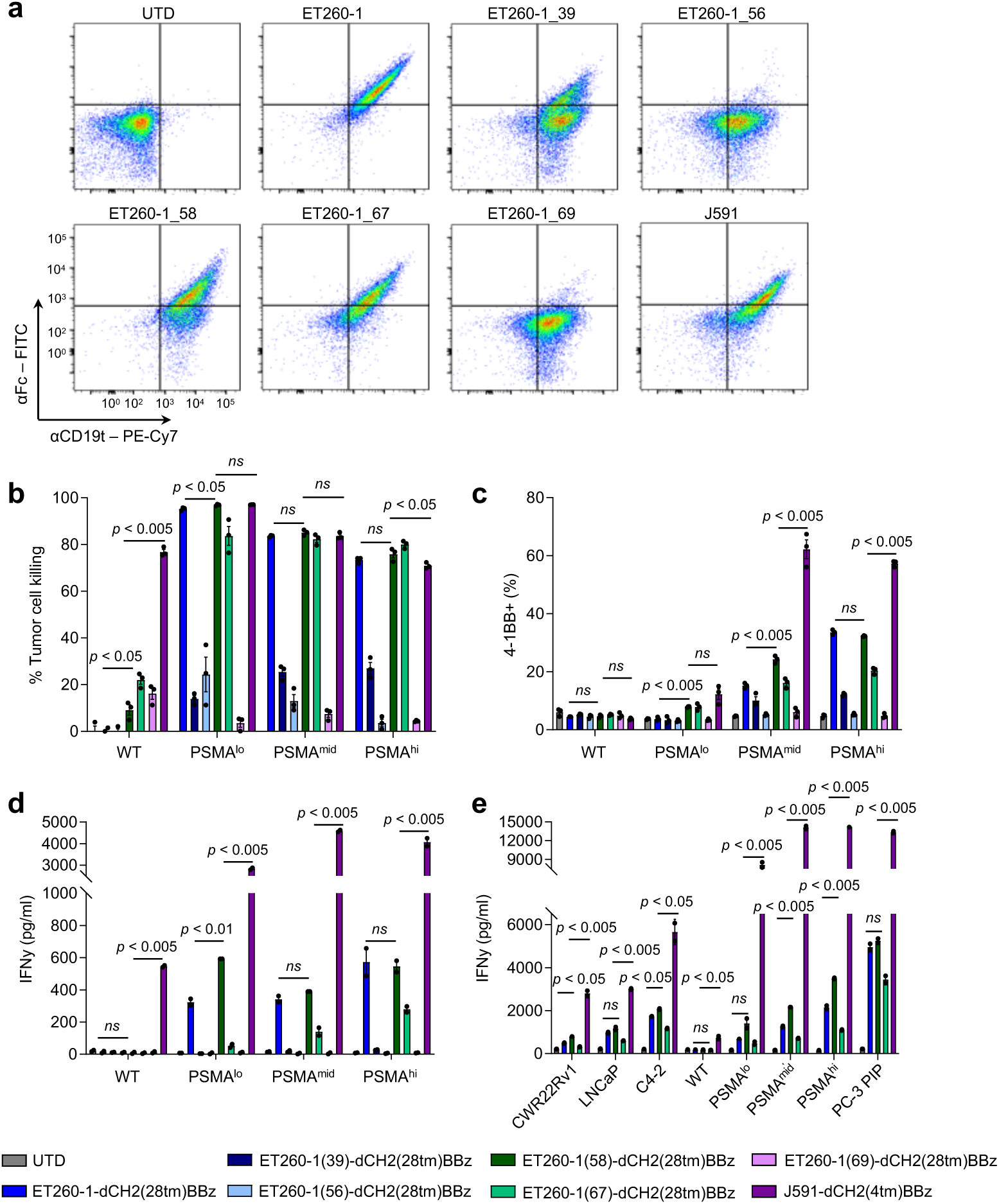
Affinity maturation of ET260-1 scFv impacts CAR stability and activity *in vitro* **a** Flow cytometric analysis of affinity matured scFvs in primary T-cells as determined by CD19t and Fc expression. **b** Tumor cell killing of AM-CARs over 72 hrs at an E:T 1:4. **c** Expression of 4-1BB on CAR T cells after 72 hr co-culture. **d** Production of IFNγ by AM-CAR T cells after 72 hrs by ELISA. **e** Activity of AM-CARs against endogenous expressers of PSMA as determined by IFNγ ELISA after a 48 hr co-culture at E:T of 1:1. Data are presented as mean values ± SEM. *P* values indicate differences between ET260-1-dCH2(28tm)BBz and ET260-1(58)-dCH2(28tm)BBz and between ET260-1(58)-dCH2(4tm)BBz and J591-dCH2(4tm)BBz using a two-tailed Student’s t-test.

**Table 1.**
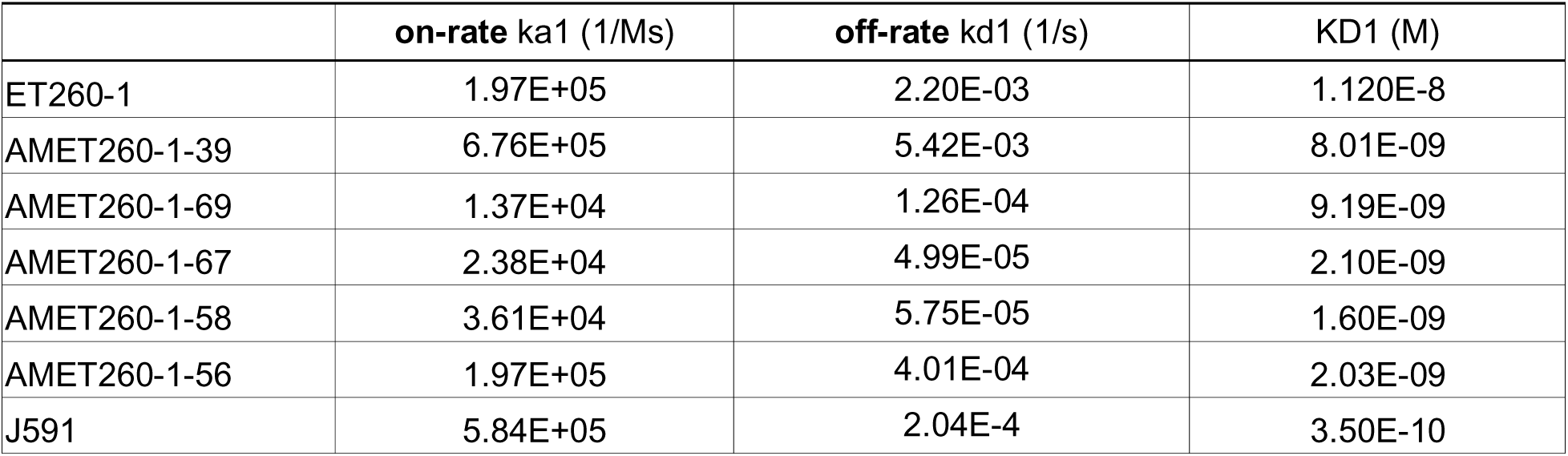
Affinities and binding kinetics of affinity-matured hPSMA scFvs. Binding to PSMA was determined by surface plasmon resonance spectroscopy (SPR). Kinetic constants from SPR-measurements were determined and KD was calculated from values of the rate constants.

Functional testing of our AM-CAR T cells showed clone 58 performed most comparably to the parental scFv when looking at tumor cell killing, activation, and IFNy after 72 hrs (**Figure 3b-d**). While we saw high tumor cell killing and IFNy secretion against PC-3 WT by J591-CAR T cells, clone 58 did not have increased activity against the WT cell line (**Figure 3b,d**). Over the course of the 8-day assay, AM-CAR T cells continued to show suboptimal activity to J591-CAR T cells (**Figure S5a,b**). We also assessed activity of our hPSMA-CAR T cells against endogenous expressers of PSMA by co-culturing T cells and quantifying IFNy production (**Figure 3e**). We saw cytokine secretion with our ET260-1 parental and AM-CAR T cells against the endogenous PSMA expressers, CWR22Rv1, LNCaP, and C4-2. Additionally, we saw that this activity was antigen-dependent with IFNy secretion increasing with antigen density, and peaking against PC-3 PIP, which has higher PSMA expression than our PC-3 PSMA^hi^ cell line (Table S2). These studies collectively suggest that in this system, there were limited improvements in CAR T cell functionality by AM of the hPSMA scFv.

### Engineering hPSMA-CAR T cells with mbIL12 rescues potent CAR T cell functionality

Previous work has demonstrated the impact of IFNy signaling on CAR T cell responses in solid tumors, including driving their direct cytolytic activity^15–17^. Since we consistently showed dampened IFNy by hPSMA-CAR T cells compared to J591-CAR T cells, associated with reduced tumor cell killing and T cell proliferation, we hypothesized that addition of IL-12 signaling, which is known to enhance IFNy secretion, may restore these activities in our hPSMA-CAR T cells. Our group recently reported on a novel membrane-bound form of IL-12 (mbIL12), which was safe and robustly improved CAR T cell functionality in solid tumors^17^. Here, we manufactured hPSMA-CAR T cells with mbIL12 to determine whether this engineering strategy could rescue potent and specific PSMA-directed CAR T cell functionality (**Figure 4a**). We transduced human T cells with both CAR (with CD19 control, ET260-1, and ET260-1(58) scFvs) and mbIL12 (**Figure 4b**). Our short-term killing assay showed comparable tumor cell killing by our hPSMA-CAR T cells with mbIL12, with no increases in non-specific targeting of PC-3 WT cells (**Figure 4c**). 4-1BB expression did not show any striking differences between the hPSMA-CARs alone and their mbIL12-expressing counterparts (**Figure 4d**). When we investigated CD25, another marker of activation, which showed higher expression in the mbIL12-containing CAR T cells compared to ET260-1-CAR T cells alone against low and high PSMA expressing tumor cells (**Figure 4e**). Similarly, IFNy secretion improved with the addition of mbIL12 to hPSMA-CAR T cells, to levels comparable to J591-CAR T cells (**Figure 4f**).

**Figure 4.**
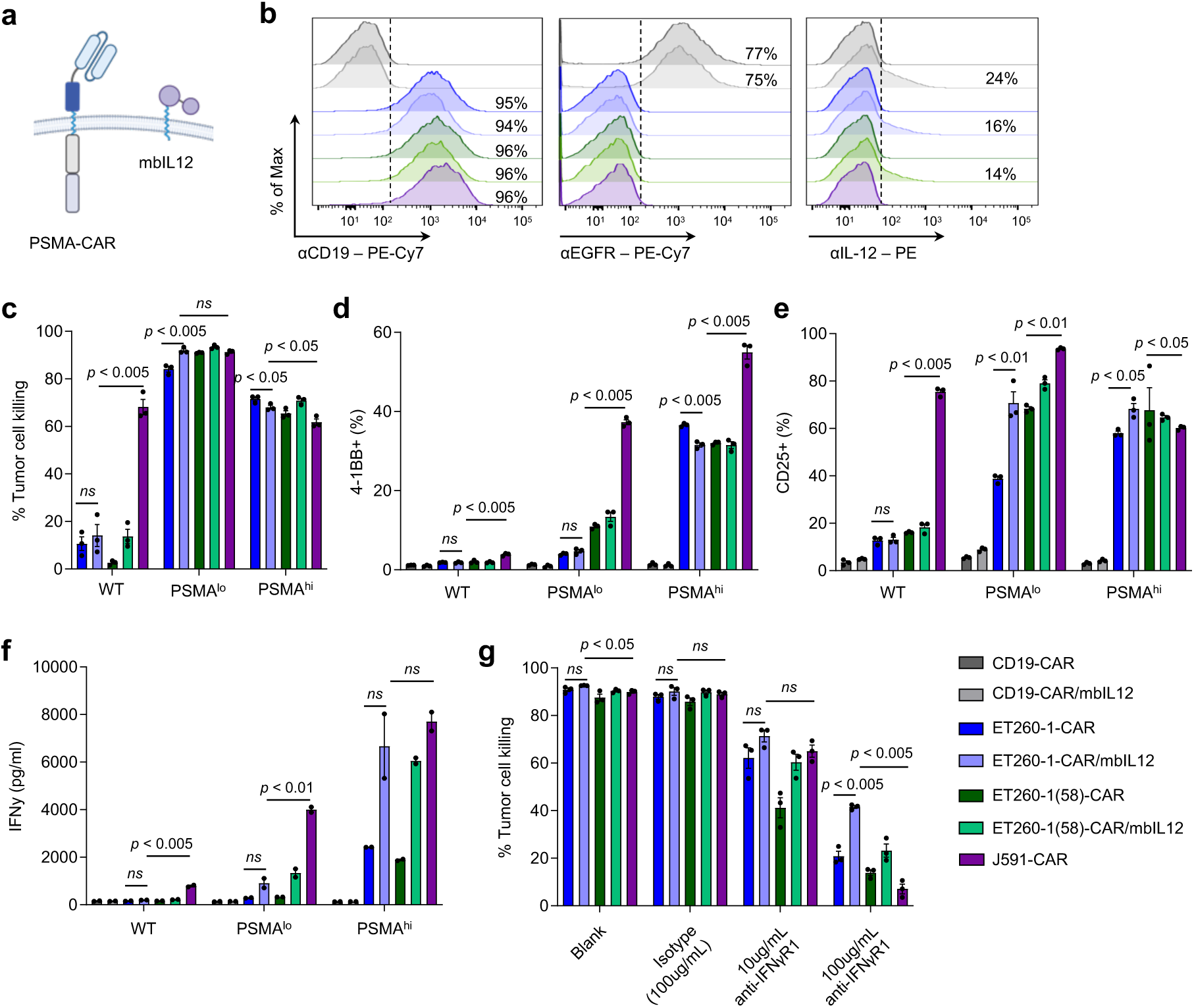
Engineering hPSMA-CARs with mbIL12 rescues CAR T cell functionality **a** Schema of hPSMA-CAR variants with mbIL12 expression. **b** Flow cytometric analysis of ET260-1 CAR and ET260-1(58) CAR with or without mbIL12 as detected by extracellular Fc and intracellular IL-12. **c** Tumor cell killing of hPSMA-CARs at an E:T of 1:4 over 72 hrs. **d** Expression of 4-1BB over 72 hrs on hPSMA-CARs. **e** Expression of CD25 on hPSMA-CARs after 72 hrs. **f** Secretion of IFNγ of hPSMA-CARs with mbIL12 after 72 hrs as determined by ELISA. **g** Tumor cell killing of hPSMA-CARs with mbIL12 and J591 against PC-3 PIP at E:T of 1:10 over 5 days at varying concentrations of anti-IFNγR1 blocking antibody and isotype control. Data are presented as mean values ± SEM. *P* values indicate differences between ET260-1-dCH2(28tm)BBz and ET260-1-dCH2(28tm)BBz/mbIL12 and between ET260-1-dCH2(28tm)BBz/mbIL12 and J591-dCH2(4tm)BBz using a two-tailed Student’s t-test.

The improved activity in the mbIL12-engineered CAR T cells to comparable or greater levels than J591-CAR T cells supported the hypothesis that this activity was at least in part IFNy-mediated. To test this further, we blocked IFNy signaling in our co-culture system, and found that tumor cell killing by J591-CAR T cells was inhibited in a dose-dependent manner with IFNyR blockade (**Figure 4g**). While tumor cell killing by hPSMA-CAR/mbIL12 T cells was also inhibited by IFNyR blockade, it was less pronounced compared with J591-CAR T cells. These results indicate that J591-CAR T cell anti-tumor activity is dependent on IFNy signaling, and that the benefits of mbIL12 on hPSMA-CAR T cells is only partially dependent on IFNy signaling and highlight IFNy-independent functionality of IL-12 signaling in our system.

The functional benefits of mbIL12 were further demonstrated in long-term killing assay (8 days), with ET260-1-CAR/mbIL12 T cells showing improved tumor cell killing compared to the parental scFv-based CAR T cells (**Figure 5a**). Importantly, T cell expansion and IFNy secretion was greater with mbIL12 compared to the parental scFv-based CAR T cells and to J591-CAR T cells (**Figure 5b-c)**. Additionally, although our ET260-1-CAR/mbIL12 T cells had improved killing against our low expresser, activity did not increase against PC-3 WT (**Figure 5a)**. Furthermore, ET260-1-CAR/mbIL12 T cells had improved expansion compared to the parental scFv against the low expresser, but expansion did not reach expansion levels seen in J591(**Figure 5b**). We also did the 8-day LTK with the AM hPSMA CAR with mbIL12 against WT and PC-3 PIP, which showed an increase in activity against PC-3 WT (**Figure S6a**). Due to the increase sensitivity to low PSMA expression and suboptimal fold expansion and cytokine secretion compared to the parental hPSMA CAR with mbIL12 (**Figure S6b,c**), we moved forward with interrogating the parental CAR with mbIL12.

**Figure 5.**
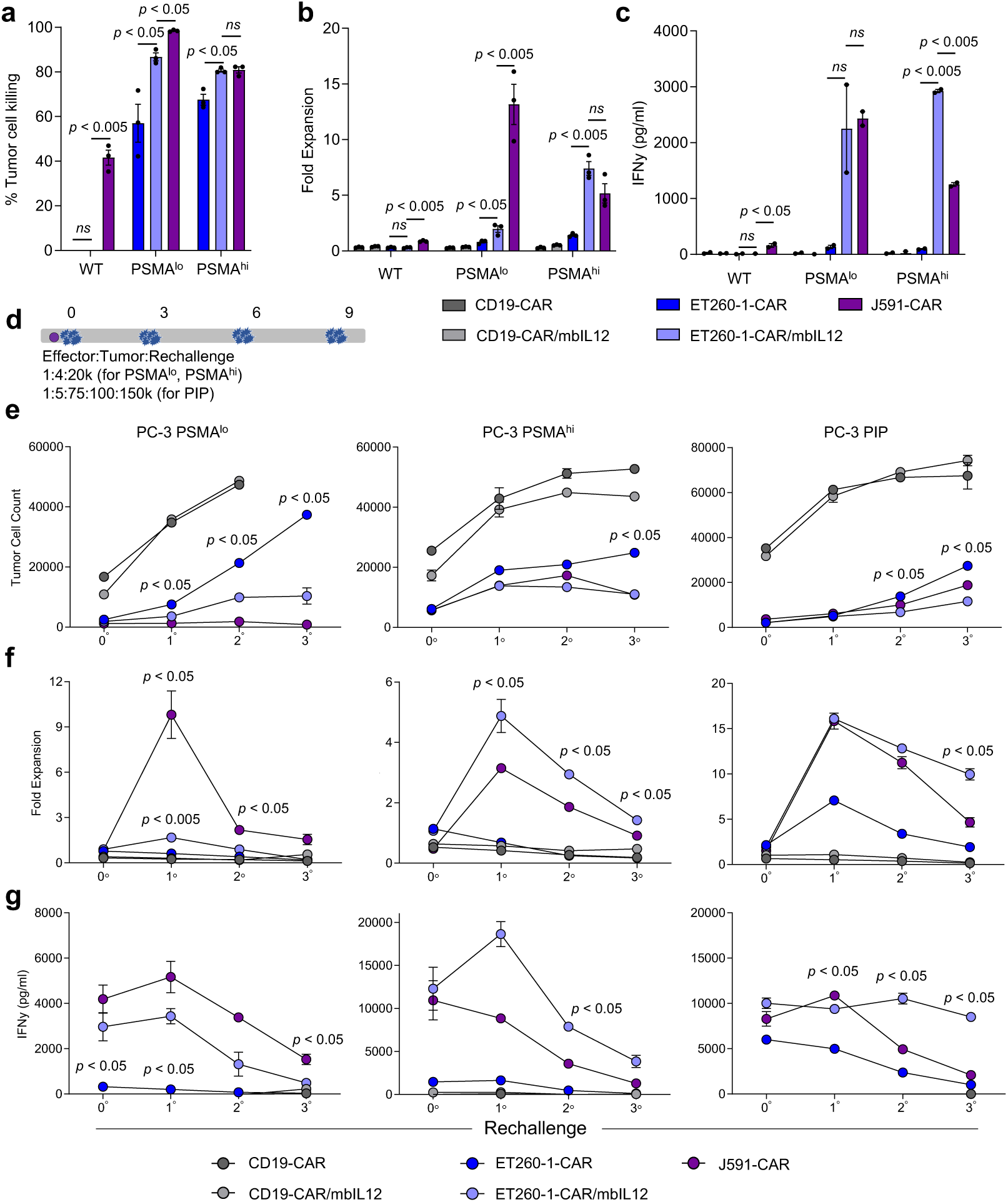
mbIL12-engineered hPSMA-CAR T cells demonstrate improved activity in a recursive tumor cell assay *in vitro* **a** Tumor cell killing of hPSMA-CARs over 8 days when co-cultured with PSMA-expressing cell lines at E:T of 1:20. **b** Fold expansion of hPSMA-CARs compared to J591-CAR after 8-day co-culture. **c** Production of IFNγ at 8 days as determined by ELISA. **d** Schema of rechallenge with PSMA-CARs co-cultured with PC-3 PSMA^lo^, PC-3 PSMA^hi^, or PC-3 PIP and rechallenged with tumor cells every 3 days. **e** PC-3 PSMA^lo^ (left), PC-3 PSMA^hi^ (center), or PC-3 PIP (right) cell counts were quantified by flow cytometry. **b** Fold expansion of T cells after each rechallenge were quantified by flow cytometry. **c** Production of IFNγ by T cells following each rechallenge as determined by ELISA. Data are presented as mean values ± SEM. *P* values indicate differences between ET260-1-dCH2(28tm)BBz and ET260-1-dCH2(28tm)BBz/mbIL12 and between ET260-1-dCH2(28tm)BBz/mbIL12 and J591-dCH2(4tm)BBz using a two-tailed Student’s t-test.

To stress test our hPSMA-CAR T cells further, we recursively challenged CAR T cells with three rechallenges of PSMA tumor cells for over the course of two weeks (**Figure 5d**). Supporting our previous *in vitro* results, ET260-1-CAR/mbIL12 T cells showed superior tumor cell killing compared to T cells expressing ET260-1-CAR alone and was comparable to J591-CAR T cells (**Figure 5e**). Additionally, ET260-1-CAR/mbIL12 T cells exhibited higher T cell expansion against PSMA high expresser and PC-3 PIP compared to both J591-CAR and ET260-1-CAR T cells alone (**Figure 5f**). Lastly, the ET260-1-CAR/mbIL12 T cells secreted higher and more sustained IFNy levels compared to J591-CAR or ET260-1-CAR T cells alone (**Figure 5g**). ET260-1-CAR/mbIL12 T cells were less sensitive to PSMA low expressing tumor cells. In contrast, J591-CAR T cells showed higher T cell expansion and IFNy secretion against PSMA low expresser. Collectively, our data supports mbIL12 engineered in improving hPSMA-CAR T cell anti-tumor activity against PSMA tumor cells, while maintaining their selectivity and specificity.

### hPSMA-CAR/mbIL12 T cells demonstrate a dampened cytokine profile in the presence of monocyte-derived suppressive M1 and M2 macrophages

Clinically evaluated PSMA-targeting strategies to date have been marred with severe adverse events attributed to cytokine-release syndrome and macrophage-activation syndrome. To evaluate the possibility of our hPSMA-CAR T cells potentiating similar toxicities, CAR T cells were cocultured in the presence of M1-polarized or M2-polarized macrophages using our previously described *in vitro* system^18^. Macrophages were differentiated and polarized into M1-like (CD80^high^, CD163^−^, CD206^low^) or M2-like (CD80^low^, CD163^+^, CD206^high^) macrophages (**Figure 6a, Figure S7**). CAR T cells, macrophages, and our PC-3 parental or PSMA^Hi^ tumor cells were cocultured at an effector:macrophage:tumor cell ratio of 1:2:4 (**Figure 6b**). As measures of activity, we quantified activation of CAR T cells via CD25 after 72 hrs and secretion of IL-6 and IFNy at 24 hrs against PC-3 WT (**Figure 6c-e**) and PC-3 PSMA^Hi^ (**Figure 6f-h**). While there was no significant difference in IL-6 secretion between our hPSMA-CAR and hPSMA CAR/mbIL12 T cells against PC-3 WT in the tumor cell only and macrophage conditions, J591-CAR T cells showed significantly higher IL-6 secretion (**Figure 6d**). We again observed significant increases in CD25 expression with the hPSMA-CAR/mbIL12 T cells compared to the hPSMA-CAR T cells alone across the different macrophage conditions (**Figure 6f**). Importantly, although there was increased activation with the hPSMA-CAR/mbIL12 T cells in the presence of M1 or M2 macrophages, there was no increases in IL-6 secretion (**Figure 6g**). In contrast, the J591-CAR T cells consistently showed higher CD25 expression and IL-6 in the tumor cell only and M2 macrophage conditions compared to hPSMA-CAR T cells (**Figure 6g**). Our hPSMA-CAR/mbIL12 T cells secreted higher IFNy in the tumor cell only condition, while the J591-CAR T cells secreted higher IFNy in the presence of M1 and M2 macrophages (**Figure 6h**). These data indicate that while hPSMA-CAR/mbIL12 T cells improve activation and IFNy production compared to the hPSMA-CAR T cells alone in the presence of M1 or M2 macrophages, the presence of mbIL12 does not increase IL-6 production, suggesting a favorable safety profile.

**Figure 6.**
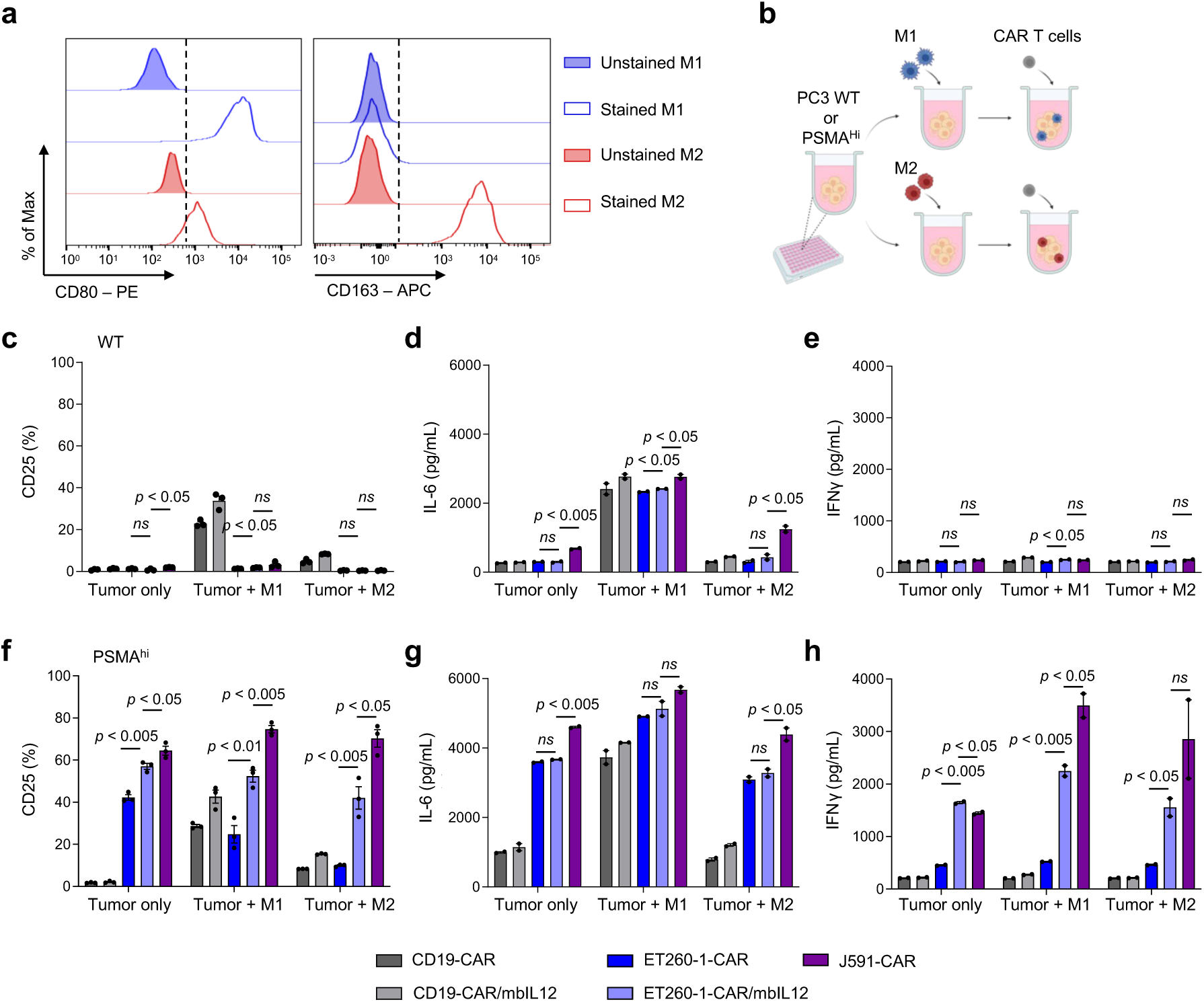
mbIL12-engineered hPSMA-CAR T cells demonstrate improved activity and cytokine profile in the presence of macrophages. **a** Confirmation of differentiation of M1-like and M2-like macrophages by detection of CD80 (left) and CD163 (right). **b** Schema of T cell suppression assay (TSA), effector: macrophage: tumor cell (E:M:T) of 1:2:4, supernatant was collected at 24 hrs and cells were processed for flow cytometry at 72 hrs. **c,f** Expression of CD25 on T cells cocultured with PC3 WT (c) or PC3-PSMA^hi^ (f) at 72 hrs. **d,g** IL-6 levels in supernatant by ELISA following coculture with PC3 WT (d) or PC3-PSMA^hi^ (g) for 24 hrs. **e,h** IFNγ levels in supernatant by ELISA following coculture with PC3 WT (e) or PC3-PSMA^hi^ (h) for 24 hrs.

### hPSMA-CAR/mbIL12 T cells demonstrate robust efficacy in a clinically relevant bone metastatic prostate cancer model

We next evaluated the therapeutic efficacy of hPSMA-CAR and hPSMA-CAR/mbIL12 T cells using an intratibial C4-2 prostate cancer bone metastasis model (**Figure 7a**). C4-2-ffluc was engrafted intratibially (i.ti.) and tumor burden was monitored using bioluminescence flux imaging. Mice were treated with a single dose of CAR T cells (CD19-CAR and CD19-CAR/mbIL12 n=5, ET260-1-CAR n=6, ET260-1-CAR/mbIL12 n=6, or J591-CAR n=6) by retro-orbital intravenous injection (i.v.). While mice treated with ET260-1-CAR T cells showed heterogeneous responses (50% complete response rate) in this model, ET260-1-CAR/mbIL12 and J591-CAR T cells showed robust and durable anti-tumor response (**Figure 7b**). While not statistically significant, mice treated with J591-CAR T cells showed 5/6 complete responses, while ET260-1-CAR/mbIL12 T cells showed 100% complete responses (**Figure 7c,d**). We observed a greater persistence of T cells in the peripheral blood of mice treated with ET260-1/mbIL12 T cells compared to J591-CAR and ET260-1-CAR T cells alone (**Figure 7e**). We validated improved therapeutic efficacy of ET260-1-CAR/mbIL12 T cells using LAPC-9, a patient-derived xenograft derived from a patient with bone metastatic prostate cancer that endogenously expresses PSMA (**Figure S8**). Mice treated with ET260-1 CAR/mbIL12 had improved survival (50% complete response) compared to the ET260-1 CAR (20% complete response) and showed no changes in body weight following treatment (**Figure S8b, c**). These results provide strong *in vivo* evidence that our ET260-1/mbIL12 CAR T cells promote robust and durable anti-tumor responses against PSMA+ prostate cancer bone metastasis.

**Figure 7.**
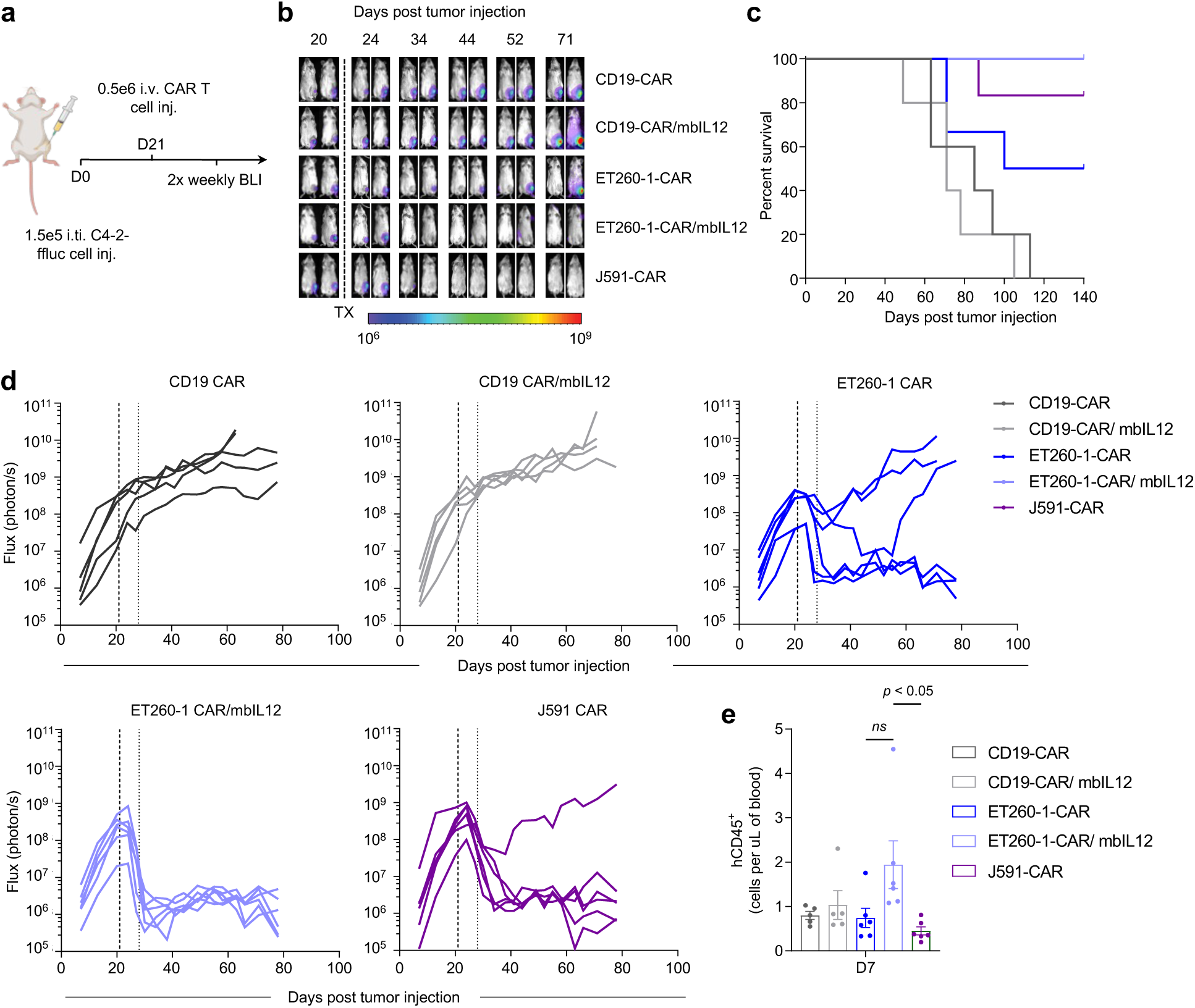
hPSMA-CAR T cells engineered with mbIL12 demonstrate robust efficacy in a clinically relevant bone metastatic prostate cancer model **a** Schema of the intratibial (i.ti.) model of C4-2_ffluc. Tumor burden was quantified using bioluminescent imaging (BLI). **b** Average tumor flux of mice. **c** Kaplan-Meier survival for CD19, CD19/mbIL12, ET260-1, ET260-1/mbIL12, and J591-CAR T cells. **D** Quantification of flux from individual mice in each treatment group (CD19 n=5, CD19/mbIL12 n=5, ET260-1 n=6, ET260-1/mbIL12 n=6, and J591 n=6). **e** Quantification of human CD45+ cells in the peripheral blood of mice treated with T cells at day 7 post-treatment. Data are presented as mean values ± SEM. *P* values indicate differences between ET260-1-dCH2(28tm)BBz/mbIL12 and J591-dCH2(4tm)BBz using a two-tailed Student’s t test.

## Discussion

Herein, we systematically optimized human PSMA-CAR T cells, which are anticipated to reduce anti-CAR immunogenicity and improve potent and selective targeting of PSMA. We first compared two human scFvs (ET260-1, ET260-2) to the humanized J591 scFv in CAR T cells. Our results highlight ET260-1 as the lead scFv over ET260-2 but with suboptimal activity compared to J591, which led us to further optimize the CAR extracellular spacer, transmembrane domains, as well as affinity mature the scFv. The combination of dCH2 and CD28tm in BBz-containing CAR T cells improved tumor cell killing, T cell activation and IFNy secretion. Although we improved upon the original CAR, we continued to observe suboptimal T cell expansion and cytokine secretion compared to J591-based CAR T cells, even following affinity-maturation of the human scFv to improve kinetics of on- and off-rates. These results demonstrate that higher scFv affinity of CARs do not necessarily result in greater functionality, but may risk greater non-specific targeting in some instances.

The proposed safety benefits of our human scFv is two-fold: 1) a human CAR avoids the potential for development of neutralizing human-anti-mouse antibodies (HAMA). HAMA has resulted in reduced persistence of CAR T cells and in some cases led to severe allergic reactions^19–22^. We have shown our ET260-1 scFv to be selective and specific to higher expression of PSMA, which we expect to mitigate potential on-target off-tumor toxicities associated with lower expression of PSMA in normal tissues including the salivary gland^23^ and in the brain^24,25^.

Studies using J591 in combination with drug conjugates, toxic payloads or as the antigen-binding domain of CAR T cells have demonstrated activity but have also been associated with unwanted toxicities^6,26^. These studies have also highlighted the tight binding of J591 to PSMA^27–30^, which our binding kinetic studies also confirmed. We postulate that J591’s activity against our PC-3 parental cell line, which has low but detectable levels of PSMA, leads to potent activation and is a likely explanation for some of the observed toxicities in PSMA-CAR T cell clinical investigations. Even when coupled with mbIL12 engineering, our hPSMA-CAR T cells demonstrated potent targeting of PSMA+ tumor cells, without increasing activation or targeting of PC3 cells with very low PSMA expression. The safety of our hPSMA-CAR T cells was further supported by lower IL-6 production in the presence of macrophages, even with mbIL12-engineering compared to J591-based CAR T cells.

Increasing evidence points to the essential role of IFNy signaling for CAR T cell activation and early killing potential against solid tumors ^15,16^. Our rationale was that by introducing an immunostimulatory cytokine to our CAR T cells we could bridge this gap in functional activity while retaining the safety and specificity of our human ET260-1 scFv-based CAR T cells. Based on recent published work from our laboratory, we combined our hPSMA-CARs with a membrane-bound IL-12 cytokine^17^. Indeed, mbIL12-engineered hPSMA-CAR T cells completely restored functional activity to similar and even greater extent than J591-CAR T cells. This activity was not entirely IFNγ-dependent, since our human PSMA-CARs still retained activity even following IFNyR blockade. Activity of the hPSMA-CAR indicates there is a mechanism of action intrinsic to our hPSMA construct that synergizes with mbIL12 for potent tumor cell killing in the absence of IFNy signaling. This also suggests there are other roles of IL12 signaling that may benefit our hPSMA-CARs. Future studies are warranted to understand the mechanisms regulating this IFNy-independent signaling by mbIL12, which may include ICAM1, FasL, or other adhesion related molecules. Importantly, this may be accomplished by similar beneficial activation using other cytokine modulators, including IL-7, IL-15, and IL-21.

We recognize that our hPSMA-CAR is limited in its single antigen targeting ability as antigen heterogeneity may limit durable response rates. As exemplified by other PSMA targeting strategies, which have transient responses^6,31,32^, and likely need recruitment of the endogenous immune system to mount a robust anti-tumor response. By incorporating mbIL12, we aim to improve the therapeutic efficacy of our hPSMA-CAR T cells while also eliciting antigen spread and endogenous anti-tumor immunity. While the current models limited our ability to investigate these phenomena, future studies using syngeneic human PSMA-knock-in (KI) mice will be of high priority.

In summary, we have optimized a human PSMA-CAR T cell therapy, which was further engineered with mbIL12 signaling, which represents an innovative approach to driving specific and potent targeting of PSMA+ cancers. These studies warrant further translational work and potential clinical testing.

## Materials and Methods

### Cell lines

Human metastatic prostate cancer cell line PC-3 (ATCC CRL-1435) was cultured in RPMI-1640 (Corning) containing 10% fetal bovine serum (FBS, Hyclone), and 1X antibiotic-antimycotic (AA, Gibco) (complete RPMI). PC-3 PIP (a kind gift from Dr. Katharina Leuckerath, UCLA) was cultured in complete RPMI. C4-2 (CRL-3314) was purchased from ATCC and cultured in DMEM:F12 (Life Technologies) with 10% FBS. C4-2 cells were transduced with lentivirus containing firefly luciferase to generate C4-2_ffluc. CWR22Rv1 (CRL-2505) (a kind gift from Dr. Robert Reiter, UCLA) and LNCaP were cultured in complete RPMI

The human fibrosarcoma cell line, HT1080 (ATCC CCL-12), and the human embryonic kidney cell line, 293T (ATCC CRL-3216), were cultured in Dulbecco’s Modified Eagles Medium (DMEM, Life Technologies) containing 10% FBS, 1X AA, 25 mM HEPES (Irvine Scientific), and 2 mM L-Glutamine (Corning™, complete DMEM).

The human prostate cancer-derived xenograft LAPC-9 (a kind gift from Dr. Robert Reiter, UCLA) was serially passaged in male NOD.Cg-Prkdcscid IL2rgtm1Wjl/SzJ (NSG) mice, and single-cell suspensions were prepared using the Miltenyi Biotec human tumor dissociation kit (cat: 130-095-929) per manufacturer’s protocol. LAPC9 cells were transduced with lentivirus to express enhanced green fluorescent protein (eGFP)/firefly luciferase (LAPC-9-eGFP-ffLuc).

### Human PSMA antibody panning

Jurkat cells engineered to over-express PSMA were used for antibody panning against Eureka Therapeutics’ E-ALPHA® phage library, a collection of human scFv antibody phage display libraries containing over 10×10^10^ unique clones. The phage libraries included naïve and semi-synthetic human scFv antibodies. The E-ALPHA® library was used to screen for the selection of human antibody constructs (e.g., scFv) specific for PSMA. In a first step, the library was negatively screened using Jurkat parental cells to eliminate clones that may recognize cell surface molecules other than PSMA. The remaining clones were screened in three rounds, using Jurkat-PSMA engineered cells to select PSMA+ clones. The positive clones were confirmed by flow cytometry for specificlly binding to hPSMA on Jurkat-PSMA cells, and not Jurkat parental cells. Finally, unique selected clones were further confirmed by flow cytometry using the PSMA-positive cancer cell lines, LnCAP, Jurkat-PSMA cells, and as negative controls, PSMA-negative cell lines Jurkat and PC3.

### DNA constructs and lentivirus production

PC-3 tumor cells were engineered to express PSMA by transduction with MSCV retrovirus carrying the human PSMA gene (Accession #:NC_000011.10). PSMA^+^ cells underwent fluorescence-activated cell sorting (FACS) using the BD FACSAria™ cell sorter. Single cell clones were generated and selected based on antigen density for low, mid, and high PSMA expression.

The ET260 scFv sequences were generated and provided by Eureka Therapeutics and described above. The J591 scFv was based on the PSMA CAR in patent US20110268656A1. The two distinct ET260 scFv chain variants with differing CDR regions were cloned into a CH2-deleted version (ΔCH2, 129-amino acid middle-length) of the IgG4 Fc spacer, CD4 transmembrane, 4-1BB intracellular costimulatory domain, and CD3ζ cytolytic signaling domain CAR construct. The 4-1BB and CD3ζ cytolytic domains were consistent throughout all future construct iterations. A T2A ribosomal skip sequence separated the CAR construct from the truncated CD19 (CD19t) tag. All CAR constructs were cloned into the epHIV7 lentiviral vector under the EF1α promoter and a GMCSFRα signal sequence. The J591 CAR was cloned into the ΔCH2 linker and CD4tm construct.

For transmembrane and linker combination comparison, a short hinge linker (HL, 22-amino acid length) was combined with a CD28 transmembrane domain (CD28tm), the ΔCH2 linker was combined with the CD28tm, and lastly a CD8 hinge (C8h, 49 amino acids) was combined with a CD8tm. All constructs contained the 4-1BB intracellular costimulatory domain with the CD3ζ signaling domain.

### Affinity maturation and clone selection

An affinity maturation library of ET260-1 scFv clones was constructed by error-prone PCR using a random mutagenesis kit (GeneMorph II, Agilent) according to the manufacturer’s instructions. Briefly, 100 ng of ET260-1 phagemids were used as template in a 20 cycle PCR reaction performed using the manufacturer’s guidelines to attain the desired mutation rate. The PCR products were purified and re-converted into phagemids. These phagemids were transduced into TG1 competent cells by electroporation; the phage library was generated by precipitation of TG1 cell culture supernatants containing phage. TG1 cells are an electrocompetent E. coli amber suppressor strain (*supE*) suitable for protein expression and preparation of antibody or peptide phage display libraries. AMET260-1 was obtained from a phage library constructed with a medium mutation rate and an effective size of 7 x 10^6^ unique clones..

Plate panning against the AMET260-1 phage library was performed using His-tagged PSMA ECD (Lys44-Ala750) protein-coated Corning® 96-well Clear Flat Bottom Polystyrene High Binding plates at a concentration of 2 ug/mL in a volume of 100 uL/well at room temperature overnight. An irrelevant His-tagged protein (MCT4) was used as a negative control. After blocking with 3% non-fat milk, three rounds of selection were carried out under more and more stringent conditions by increasing the incubation and washing times. The washing buffer contained 5 ug/mL of free PSMA ECD protein to compete with plate-coated antigen.

Affinity-matured (AM) scFvs were cloned into the ΔCH2/CD28tm vector. The membrane-bound IL-12 (mbIL12) construct was generated using the p35 and p40 genes (p35, NC_000003.12; p40, NC_000005.10) separated by a G4S spacer, and linked to the CD28 transmembrane domain as described previously^17^.

Lentivirus was generated as previously described^33^. Lentiviral titers, as determined by CD19t, EGFRt or IL12 expression, were quantified using HT1080 cells.

#### SPR

The binding affinity of the scFv clones to the extracellular domain of human PMSA (Lys44-Ala750) was measured by Surface Plasmon Resonance using a BiaCore^TM^ X-100 instrument (Cytiva). The binding parameters between the anti-PMSA scFvs and PSMA protein were measured using a Protein G Sensor Chip (Cytiva) according to the manufacturer’s instructions for Multi-Cycle Kinetics (MCK) analysis. Briefly, the recombinant N-terminal Fc-tagged PSMA protein (hFc-PMSA) was loaded onto the protein G Sensor Chip at 10 µg/mL. The scFv proteins were passed over the sensor chip at varying concentrations of 10, 5, 2.5, 1.25, 0.625, 0.3125, 0 µg/mL in consecutive runs. The data was analyzed using a 1:1 binding site mode and BiaCoreTM X-100 Evaluation Software. The binding parameters, association (on-rate) constant *k*a, dissociation (off-rate) constant *k*d, and equilibrium dissociation constant KD were calculated.

### T cell isolation, lentiviral transduction, and ex vivo expansion

Leukapheresis products were obtained from consented research participants (healthy donors) under protocols approved by the City of Hope (COH) Internal Review Board (IRB). On the day of leukapheresis, peripheral blood mononuclear cells (PBMC) were isolated by using SepMate™ PBMC Isolation Tubes (StemCell Technologies) followed by multiple washes in PBS with 2% FBS. To obtain depleted PBMC cells (dPBMC), cells were depleted of CD25 and CD14 using CD25 and CD14 beads (Miltenyi Biotec) and sorted using autoMACS® Pro Separator. Depletion was confirmed by flow cytometry and cells were frozen for future use in CryoStor® CS5 cryopreservation media (BioLife Solutions).

T cell activation and transduction were performed as previously described^33^. Where indicated, cells underwent a second lentiviral transduction 24 hrs after the first transduction. Cells were replenished with fresh cytokine X-VIVO mixture every 2-3 days. Cells were *ex vivo* manufactured, enriched for CD19t or EGFRt, and cryopreserved as previously described. CAR T cells were characterized pre- and post-enrichment for purity and phenotype by flow cytometry.

For in vitro assays, cells were thawed one day prior and rested overnight. Cells were resuspended in 1x cytokine X-VIVO at 1 million cells per mL. For in vivo studies cells were thawed on the day of treatment and resuspended at desired treatment concentration in 1x PBS.

### Flow cytometry

For flow cytometric analysis, cells were processed in FACS staining solution (FSS) (Hank’s balanced salt solution without Ca2+, Mg2+, or phenol red (HBSS-/-, Life Technologies) containing 2% FBS and 0.5g sodium azide in 500mL). Cells were washed once followed by incubation with antibodies for 30 min at 4° in the dark. Antibodies were conjugated with Brilliant Violet 510 (BV510), Brilliant Violet 570 (BV570), SuperBright 600 (SB600), Brilliant Violet 650 (BV650), fluorescein isothiocyanate (FITC), phycoerythrin (PE), peridinin chlorophyll protein complex (PerCP), PE-CF594, PE-Cy5, PerCP-Cy5.5, PerCP-eFlour710, PECy7, allophycocyanin (APC), or APC-Cy7 (or APC-eFluor780), Alexa Fluor 700 (AF700).

Antibodies against human antigens used include: CD3 (BD Biosciences, Cat: 563109, Clone: SK7), CD4 (BD BioLegend, Cat: 300534, Clone: RPA-T4), CD8 (Invitrogen, Cat: 56-008-742, Clone: SK1), CD19 (BD Biosciences, Cat: 557835, Clone: SJ25C1), CD45 (BD Biosciences, Cat: 563204, Clone: HI30), CD69 (BD Biosciences, Cat: 341652, Clone: L78), CD137 (Invitrogen, Cat: 63-137-942, Clone: 4B4-1), Tim-3 (BioLegend, Cat: 345028, Clone: F38-2E2), IL-12 (BD Bioscience, Cat: 554575, Clone: C11.5), Lag-3 (BD Biosciences, Cat: 565718, Clone: T47-530), PD-L1 (Invitrogen, Cat: 15-598-342, Clone: MIH1), CD25 (Invitrogen, Cat: 46-0259-42, Clone: BC96), PSMA (BioLegend, Cat: 342508, Clone: LNI-17), PD-1 (eBiosciences, Cat: 47-2799-42, Clone: J105).

Cell viability was determined using 4′, 6-diamidino-2-phenylindole (DAPI, Sigma, Cat: D8417). Unless otherwise stated, antibodies were used at a dilution of 1:100. Flow cytometry was performed on a MACSQuant Analyzer 10 or MACSQuant Analyzer 16 (Miltenyi Biotec), and the data were analyzed with FlowJo software (v10.8.1, TreeStar).

For intracellular flow cytometry, cells were processed using the BD Cytofix/Cytoperm fixation permeabilization kit (Cat: 554714) according to the manufacturer’s protocol.

PSMA quantification was done using Bangs Laboratories Quantum™ Simply Cellular® mouse IgG1 kit per manufacturer’s instructions.

### In vitro macrophage differentiation

Primary human M1-like and M2-like macrophages were differentiated as previously described^18^. Briefly, frozen human monocytes were thawed and cultured in cytokine-containing RPMI + 10% FBS and differentiated for 7–10 days. To differentiate M1-like macrophages, cells were cultured with GM-CSF (BioLegend, 572903). The media was changed once after 3–5 days to media containing GM-CSF, IFN-γ (BioLegend, 570202), LPS (Sigma-Aldrich, L3012-5MG) and IL-6 (BioLegend, 570804). To differentiate M2-like macrophages, cells were cultured with M-CSF (BioLegend, Cat: 574804). The media was changed once after 3–5 days to media containing M-CSF, IL-4 (BioLegend, 574004), IL-13 (BioLegend, 571102) and IL-6. All cytokines and LPS were used at 20 ng/mL. After differentiation, macrophages were lifted using PBS-EDTA, and phenotype was assessed by flow cytometry to confirm successful polarization. Cells were counted and used for further studies.

### In vitro cell killing

For tumor cell killing assays, T cells and tumor cells were co-cultured at indicated effector: tumor (E:T) ratios in X-VIVO with 10% FBS in 96-well plates. Co-cultures set up for 72 hr processing timepoints, an E:T of 1:4 was used and for 8 day cocultures an E:T of 1:20 was used.Cells were processed for flow cytometry at indicated timepoints. CAR T cell killing was calculated by comparing cell counts relative to counts of target cells co-cultured with untransduced T cells (UTD) or non-targeting CARs. Tumor cells were gated as DAPI-negative, single cells, CD45-negative.

For recursive tumor cell challenge assays, T cells were initially co-cultured with PSMA expressing cell lines at 1:4 E:T for the PC-3 PSMA^hi/lo^ and 1:5 E:T for PC-3 PIP. CAR T cells co-cultured with PC-3 PSMA^hi/lo^ were rechallenged every 3 days up to 3 times with twenty-thousand tumor cells. CAR T cells co-cultured with PC-3 PIP were rechallenged every 3 days, first with 75k tumor cells, then 100k tumor cells, and the final rechallenge with 150k tumors cells. This was due to the high proliferation of the CAR T cells and to ensure an adequate rechallenge. The remaining viable cells were quantified by flow cytometry to determine tumor cell and T cell counts prior to every rechallenge, and 3 days after the last rechallenge.

### In vitro T cell suppression assay (TSA)

CAR T cells, macrophages, and target tumors were co-cultured in X-Vivo + 10% FBS in the absence of exogenous cytokines in 96-well plates. Cells were plated at an effector:macrophage:target ratio of 1:2:4 to model prostate cancer with PC3 WT or PC3-PSMA^hi^ cells. Supernatant was collected after 24 hrs and 72 hrs for ELISA, and cells were trypsinized and collected for flow cytometry after 72 hrs. Tumor cell killing and T cell activation were evaluated after 72 hrs by flow cytometry.

### ELISA cytokine assays

Supernatant collected from tumor cell killing assays were collected at indicated timepoints and stored at -20°C for future use. Supernatants were used to analyze secretion of IFNγ with the ELISA Ready-SET-GO!® kit (Cat: 88-7316-88) according to manufacturer’s protocol. Plates were read at 450nm and 650nm using Cytation5 imaging reader with Gen5 microplate software V3.08 (BioTek).

### In vivo animal studies

All animal experiments were performed under protocols approved by the City of Hope Institutional Animal Care and Use Committee. All mice were co-housed in a maximum barrier, pathogen- and opportunist-free animal facility. For intratibial (i.ti.) tumor studies, C4-2 (0.15×10^6^) and LAPC-9 (0.75 x10^5^) were prepared in a final volume of 30 µL HBSS−/− and injected in the tibia. Tumor growth was monitored at 1-2x a week via non-invasive bioluminescence imaging (Xenogen, LagoX) and flux signals were analyzed with Aura software (Spectral Instruments Imaging). To image mice, 150 µL of D-luciferin potassium salt (Perkin-Elmer) in PBS at 4.29 mg/mouse was injected intra intraperitoneally (i.p.). Mice were treated with T cells (1×10^6^ or 0.5×10^6^) in 100µL PBS retro-orbital (i.v.r.o) on day 15 or day 21. Humane endpoints were used in determining survival. At pre-determined time points or at moribund status, mice were euthanized. Mice were euthanized by CO2 and cervical dislocation upon signs of distress, such as labored or difficulty breathing, apparent weight loss, impaired mobility, or evidence of being moribund.

### Statistical analysis and reproducibility

Data are presented as mean ± standard error of mean (SEM), unless otherwise stated. Statistical comparisons between groups were performed using the unpaired two-tailed Student’s t-test to calculate p-value, unless otherwise stated.

## Acknowledgments

We thank the staff members of the following cores at the Beckman Research Institute at City of Hope Comprehensive Cancer Center: Animal Facility and Small Animal Imaging for their excellent technical assistance. Work performed in the Small Animal Imaging Core was supported by the National Cancer Institute of the National Institutes of Health under grant number P30CA033572. Research reported in this publication was supported by Eureka Therapeutics Sponsored Research Agreement (PI: Priceman), the Fiterman Family Foundation fund and the Doug and Rhonda Collier Foundation fund.

The content is solely the responsibility of the authors and does not necessarily represent the official views of the National Institutes of Health.

## Author contributions

S.J.P., along with L.S.L. provided the conception and construction of the study. L.S.L., Z.C., Y.Y., J.P.M., Z.Y., K.Z., J.Y., W-C.C. and S.J.P. provided the design of experimental procedures, data analysis, and/or interpretation. L.S.L., Z.C., Y.Y., J.P.M., Z.Y., K.Z., J.Y., W-C.C. performed experiments. L.S.L., S.J.P. wrote the manuscript. Z.C., S.J.F., and V.W.C. assisted in writing/editing the manuscript. S.J.P. supervised the study. All authors reviewed the manuscript.

## Data Availability Statement

All data associated with this study are present in the paper or the Supplemental information. Materials are available from S.J.P. under a material transfer agreement.

## Supplemental Titles

Figure S1. Confirmation and Characterization of unique panning clones positively bind to PSMA.

**a.** Flow cytometric analysis of the PSMA-expression positive cells, LnCAP, Jurkat - PSMA, and the PSMA negative cells, Jurkat and PC3 cells, stained with each clone of phage and followed by PE-anti-M13 mAb, respectively. **b.** Kinetics of two unique clones in BiTE format toward hFc tagged ECD of PSMA_44-750_ on a protein G sensor chip. The kinetic lines from top to bottom represent 10, 3.33, 1.11, 0.37, 0.123 ug/ml.

Figure S2. hPSMA-CARs are efficiently transduced and enriched in primary T cells

**a.** Flow cytometric analysis of scFv variants determined by the truncated CD19 tag (CD19t). **b.** Flow cytometric analysis of ET260-1 extracellular linker and transmembrane domain variants determined by CD19t.

Figure S3. Quantification of PSMA on surface of prostate tumor cell lines

**a.** Histograms plots of PSMA expression on prostate cancer cell lines used. **b.** Quantification of PSMA expression using the Bangs Laboratories Quantum™ Simply Cellular® mouse IgG1 kit.

Figure S4. Binding kinetics of selected AM clones in scFv format

Surface plasmon resonance (SPR) sensorgrams of AM clones with the kinetic lines from top to bottom representing 10, 5, 2.5, 1.25, 0.625, 0.3125 ug/ml of scFv.

Figure S5. Functional assessment of AM-CAR T cells in 8-day LTK

**a.** Tumor cell killing of hPSMA-CARs over 72 hrs when co-cultured with PSMA-expressing cell lines at E:T of 1:8. **b.** Production of IFNγ at 8 days as determined by ELISA.

Figure S6. Functional assessment of hPSMA-CAR T cells engineered with mbIL12 in 8-day LTK

**a.** Tumor cell killing of hPSMA-CARs over 8 days when co-cultured with PC-3 PIP at E:T of 1:20. **b.** Fold expansion of hPSMA-CARs compared to J591-CAR after 8-day co-culture. **c** Production of IFNγ at 8 days as determined by ELISA.

Figure S7. Phenotype of M1-like and M2-like macrophages following *in vitro* polarization

**a.** Confirmation of differentiation of M1-like and M2-like macrophages by detection of CD206.

Figure S8. hPSMA-CAR T cells engineered with mbIL12 demonstrate efficacy against LAPC-9 bone metastasis xenograft model

**a.** Schema of the intratibial (i.ti.) model of LAPC-9_eGFP-FFluc. Tumor burden was quantified using bioluminescent imaging (BLI). **b.** Kaplan-Meier survival for CD19-CAR, CD19-CAR/mbIL12, ET260-1-CAR, and ET260-1-CAR/mbIL12 T cell-treated mice. **c.** Weight change of mice relative to initial weight prior to T cell treatment. **d,e.** Quantification of flux from the tibia **d.** and whole body **e.** from individual mice in each treatment group (CD19-CAR; n=10, CD19-CAR/mbIL12; n=10, ET260-1-CAR; n=10, ET260-1-CAR/mbIL12; n=10)

Table S1. Binding kinetics of parental scFvs against PSMA by SPR

Binding to PSMA was determined by surface plasmon resonance spectroscopy (SPR). Constants from SPR-measurements were determined and KD was calculated from values of the rate constants.

Table S2. Quantification of PSMA on the surface of prostate tumor cell lines

Density of PSMA was determined using Bangs Laboratories, Inc Simply Cellular IgG quantification kit per manufacturer’s protocols.

Table S3. Binding kinetics of affinity matured scFvs against PSMA by SPR

Binding to PSMA was determined by surface plasmon resonance spectroscopy (SPR) for the 10 selected AM clones. Kinetic constants from SPR-measurements were determined and KD was calculated from values of the rate constants.

## References

1. Siegel, R.L., Miller, K.D., Wagle, N.S., and Jemal, A. (2023). Cancer statistics, 2023. CA: a cancer journal for clinicians 73, 17-48. 10.3322/caac.21763.

2. Cai, M., Song, X.-L., Li, X.-A., Chen, M., Guo, J., Yang, D.-H., Chen, Z., and Zhao, S.-C. (2023). Current therapy and drug resistance in metastatic castration-resistant prostate cancer. Drug Resistance Updates 68, 100962. 10.1016/j.drup.2023.100962.

3. Alzubi, J., Dettmer-Monaco, V., Kuehle, J., Thorausch, N., Seidl, M., Taromi, S., Schamel, W., Zeiser, R., Abken, H., Cathomen, T., and Wolf, P. (2020). PSMA-Directed CAR T Cells Combined with Low-Dose Docetaxel Treatment Induce Tumor Regression in a Prostate Cancer Xenograft Model. Molecular Therapy - Oncolytics 18, 226–235. 10.1016/j.omto.2020.06.014.

4. Friedrich, M., Raum, T., Lutterbuese, R., Voelkel, M., Deegen, P., Rau, D., Kischel, R., Hoffmann, P., Brandl, C., Schuhmacher, J., et al. (2012). Regression of human prostate cancer xenografts in mice by AMG 212/BAY2010112, a novel PSMA/CD3-Bispecific BiTE antibody cross-reactive with non-human primate antigens. Mol Cancer Ther 11, 2664–2673. 10.1158/1535-7163.Mct-12-0042.

5. Dorff, T.B., Blanchard, S., Martirosyan, H., Adkins, L., Dhapola, G., Shishido, S., Wagner, J.R., Stiller, T., D’Apuzzo, M., Kuhn, P., et al. (2023). Final results from phase I study of PSCA-targeted chimeric antigen receptor (CAR) T cells in patients with metastatic castration resistant prostate cancer (mCRPC). Journal of Clinical Oncology 41, 5019–5019. 10.1200/JCO.2023.41.16_suppl.5019.

6. Narayan, V., Barber-Rotenberg, J.S., Jung, I.-Y., Lacey, S.F., Rech, A.J., Davis, M.M., Hwang, W.-T., Lal, P., Carpenter, E.L., Maude, S.L., et al. (2022). PSMA-targeting TGFβ-insensitive armored CAR TLcells in metastatic castration-resistant prostate cancer: a phase 1 trial. Nature Medicine 28, 724–734. 10.1038/s41591-022-01726-1.

7. Deegen, P., Thomas, O., Nolan-Stevaux, O., Li, S., Wahl, J., Bogner, P., Aeffner, F., Friedrich, M., Liao, M.Z., Matthes, K., et al. (2021). The PSMA-targeting Half-life Extended BiTE Therapy AMG 160 has Potent Antitumor Activity in Preclinical Models of Metastatic Castration-resistant Prostate Cancer. Clinical cancer research : an official journal of the American Association for Cancer Research 27, 2928–2937. 10.1158/1078-0432.Ccr-20-3725.

8. Giraudet, A.L., Kryza, D., Hofman, M., Moreau, A., Fizazi, K., Flechon, A., Hicks, R.J., and Tran, B. (2021). PSMA targeting in metastatic castration-resistant prostate cancer: where are we and where are we going? Therapeutic advances in medical oncology 13, 17588359211053898. 10.1177/17588359211053898.

9. Bailis, J., Deegen, P., Thomas, O., Bogner, P., Wahl, J., Liao, M., Li, S., Matthes, K., Nägele, V., Rau, D., et al. (2019). Preclinical evaluation of AMG 160, a next-generation bispecific T cell engager (BiTE) targeting the prostate-specific membrane antigen PSMA for metastatic castration-resistant prostate cancer (mCRPC). Journal of Clinical Oncology 37, 301–301. 10.1200/JCO.2019.37.7_suppl.301.

10. Skokos, D., Waite, J.C., Haber, L., Crawford, A., Hermann, A., Ullman, E., Slim, R., Godin, S., Ajithdoss, D., Ye, X., et al. (2020). A class of costimulatory CD28-bispecific antibodies that enhance the antitumor activity of CD3-bispecific antibodies. Science Translational Medicine 12, eaaw7888. 10.1126/scitranslmed.aaw7888.

11. Klein Nulent, T.J.W., Valstar, M.H., de Keizer, B., Willems, S.M., Smit, L.A., Al-Mamgani, A., Smeele, L.E., van Es, R.J.J., de Bree, R., and Vogel, W.V. (2018). Physiologic distribution of PSMA-ligand in salivary glands and seromucous glands of the head and neck on PET/CT. Oral Surg Oral Med Oral Pathol Oral Radiol 125, 478–486. 10.1016/j.oooo.2018.01.011.

12. Silver, D.A., Pellicer, I., Fair, W.R., Heston, W.D., and Cordon-Cardo, C. (1997). Prostate-specific membrane antigen expression in normal and malignant human tissues. Clinical cancer research : an official journal of the American Association for Cancer Research 3, 81–85.

13. Emperumal, C.P., Villa, A., Hwang, C., Oh, D., Fong, L., Aggarwal, R., and Keenan, B.P. (2024). Oral Toxicities of PSMA-Targeted Immunotherapies for The Management of Prostate Cancer. Clinical Genitourinary Cancer 22, 380–384. 10.1016/j.clgc.2023.12.008.

14. Laidler, P., Dulińska, J., Lekka, M., and Lekki, J. (2005). Expression of prostate specific membrane antigen in androgen-independent prostate cancer cell line PC-3. Archives of Biochemistry and Biophysics 435, 1–14. 10.1016/j.abb.2004.12.003.

15. Alizadeh, D., Wong, R.A., Gholamin, S., Maker, M., Aftabizadeh, M., Yang, X., Pecoraro, J.R., Jeppson, J.D., Wang, D., Aguilar, B., et al. (2021). IFNγ Is Critical for CAR T Cell–Mediated Myeloid Activation and Induction of Endogenous Immunity. Cancer Discovery 11, 2248–2265. 10.1158/2159-8290.CD-20-1661.

16. Larson, R.C., Kann, M.C., Bailey, S.R., Haradhvala, N.J., Llopis, P.M., Bouffard, A.A., Scarfó, I., Leick, M.B., Grauwet, K., Berger, T.R., et al. (2022). CAR T cell killing requires the IFNγR pathway in solid but not liquid tumours. Nature 604, 563–570. 10.1038/s41586-022-04585-5.

17. Lee, E.H.J., Murad, J.P., Christian, L., Gibson, J., Yamaguchi, Y., Cullen, C., Gumber, D., Park, A.K., Young, C., Monroy, I., et al. (2023). Antigen-dependent IL-12 signaling in CAR T cells promotes regional to systemic disease targeting. Nature Communications 14, 4737. 10.1038/s41467-023-40115-1.

18. Yamaguchi, Y., Gibson, J., Ou, K., Lopez, L.S., Ng, R.H., Leggett, N., Jonsson, V.D., Zarif, J.C., Lee, P.P., Wang, X., et al. (2022). PD-L1 blockade restores CAR T cell activity through IFN-γ-regulation of CD163+ M2 macrophages. Journal for immunotherapy of cancer 10. 10.1136/jitc-2021-004400.

19. Zhang, C., Wang, L., Zhang, Q., Shen, J., Huang, X., Wang, M., Huang, Y., Chen, J., Xu, Y., Zhao, W., et al. (2023). Screening and characterization of the scFv for chimeric antigen receptor T cells targeting CEA-positive carcinoma. Frontiers in immunology 14, 1182409. 10.3389/fimmu.2023.1182409.

20. Maus, M.V., Haas, A.R., Beatty, G.L., Albelda, S.M., Levine, B.L., Liu, X., Zhao, Y., Kalos, M., and June, C.H. (2013). T Cells Expressing Chimeric Antigen Receptors Can Cause Anaphylaxis in Humans. Cancer Immunology Research 1, 26–31. 10.1158/2326-6066.CIR-13-0006.

21. Till, B.G., Jensen, M.C., Wang, J., Chen, E.Y., Wood, B.L., Greisman, H.A., Qian, X., James, S.E., Raubitschek, A., Forman, S.J., et al. (2008). Adoptive immunotherapy for indolent non-Hodgkin lymphoma and mantle cell lymphoma using genetically modified autologous CD20-specific T cells. Blood 112, 2261–2271. 10.1182/blood-2007-12-128843.

22. Blanco, I., Kawatsu, R., Harrison, K., Leichner, P., Augustine, S., Baranowska-Kortylewicz, J., Tempero, M., and Colcher, D. (1997). Antiidiotypic response against murine monoclonal antibodies reactive with tumor-associated antigen TAG-72. J Clin Immunol 17, 96–106. 10.1023/a:1027396714623.

23. Israeli, R.S., Powell, C.T., Corr, J.G., Fair, W.R., and Heston, W.D. (1994). Expression of the prostate-specific membrane antigen. Cancer research 54, 1807–1811.

24. Kunikowska, J., Czepczyński, R., Pawlak, D., Koziara, H., Pełka, K., and Królicki, L. (2022). Expression of glutamate carboxypeptidase II in the glial tumor recurrence evaluated in vivo using radionuclide imaging. Scientific Reports 12, 652. 10.1038/s41598-021-04613-w.

25. Rischpler, C., Beck, T.I., Okamoto, S., Schlitter, A.M., Knorr, K., Schwaiger, M., Gschwend, J., Maurer, T., Meyer, P.T., and Eiber, M. (2018). 68-Ga-PSMA-HBED-CC Uptake in Cervical, Celiac, and Sacral Ganglia as an Important Pitfall in Prostate Cancer PET Imaging. Journal of Nuclear Medicine 59, 1406–1411. 10.2967/jnumed.117.204677.

26. Tagawa, S.T., Beltran, H., Vallabhajosula, S., Goldsmith, S.J., Osborne, J., Matulich, D., Petrillo, K., Parmar, S., Nanus, D.M., and Bander, N.H. (2010). Anti–prostate-Specific membrane antigen-based radioimmunotherapy for prostate cancer. Cancer 116, 1075–1083. 10.1002/cncr.24795.

27. Smith-Jones, P.M., Vallabahajosula, S., Goldsmith, S.J., Navarro, V., Hunter, C.J., Bastidas, D., and Bander, N.H. (2000). In Vitro Characterization of Radiolabeled Monoclonal Antibodies Specific for the Extracellular Domain of Prostate-specific Membrane Antigen1. Cancer research 60, 5237–5243.

28. Fracasso, G., Bellisola, G., Cingarlini, S., Castelletti, D., Prayer-Galetti, T., Pagano, F., Tridente, G., and Colombatti, M. (2002). Anti-tumor effects of toxins targeted to the prostate specific membrane antigen. The Prostate 53, 9–23. 10.1002/pros.10117.

29. McDevitt, M.R., Barendswaard, E., Ma, D., Lai, L., Curcio, M.J., Sgouros, G., Ballangrud, A.s.M., Yang, W.-H., Finn, R.D., Pellegrini, V., et al. (2000). An α-Particle Emitting Antibody ([213Bi]J591) for Radioimmunotherapy of Prostate Cancer1. Cancer research 60, 6095–6100.

30. Frigerio, B., Fracasso, G., Luison, E., Cingarlini, S., Mortarino, M., Coliva, A., Seregni, E., Bombardieri, E., Zuccolotto, G., Rosato, A., et al. (2013). A single-chain fragment against prostate specific membrane antigen as a tool to build theranostic reagents for prostate cancer. European Journal of Cancer 49, 2223–2232. 10.1016/j.ejca.2013.01.024.

31. Sartor, O., de Bono, J., Chi, K.N., Fizazi, K., Herrmann, K., Rahbar, K., Tagawa, S.T., Nordquist, L.T., Vaishampayan, N., El-Haddad, G., et al. (2021). Lutetium-177–PSMA-617 for Metastatic Castration-Resistant Prostate Cancer. New England Journal of Medicine 385, 1091–1103. 10.1056/NEJMoa2107322.

32. Stein, M.N., Zhang, J., Kelly, W.K., Wise, D.R., Tsao, K., Carneiro, B.A., Falchook, G.S., Sun, F., Govindraj, S., Sims, J.S., et al. (2023). Preliminary results from a phase 1/2 study of co-stimulatory bispecific PSMAxCD28 antibody REGN5678 in patients (pts) with metastatic castration-resistant prostate cancer (mCRPC). Journal of Clinical Oncology 41, 154–154. 10.1200/JCO.2023.41.6_suppl.154.

33. Priceman, S.J., Gerdts, E.A., Tilakawardane, D., Kennewick, K.T., Murad, J.P., Park, A.K., Jeang, B., Yamaguchi, Y., Yang, X., Urak, R., et al. (2018). Co-stimulatory signaling determines tumor antigen sensitivity and persistence of CAR T cells targeting PSCA+ metastatic prostate cancer. Oncoimmunology 7, e1380764. 10.1080/2162402x.2017.1380764.

