## Supplemental Figures for "IL12-engineered human PSMA-CAR T cells for the treatment of advanced prostate cancer"

### Figure S1

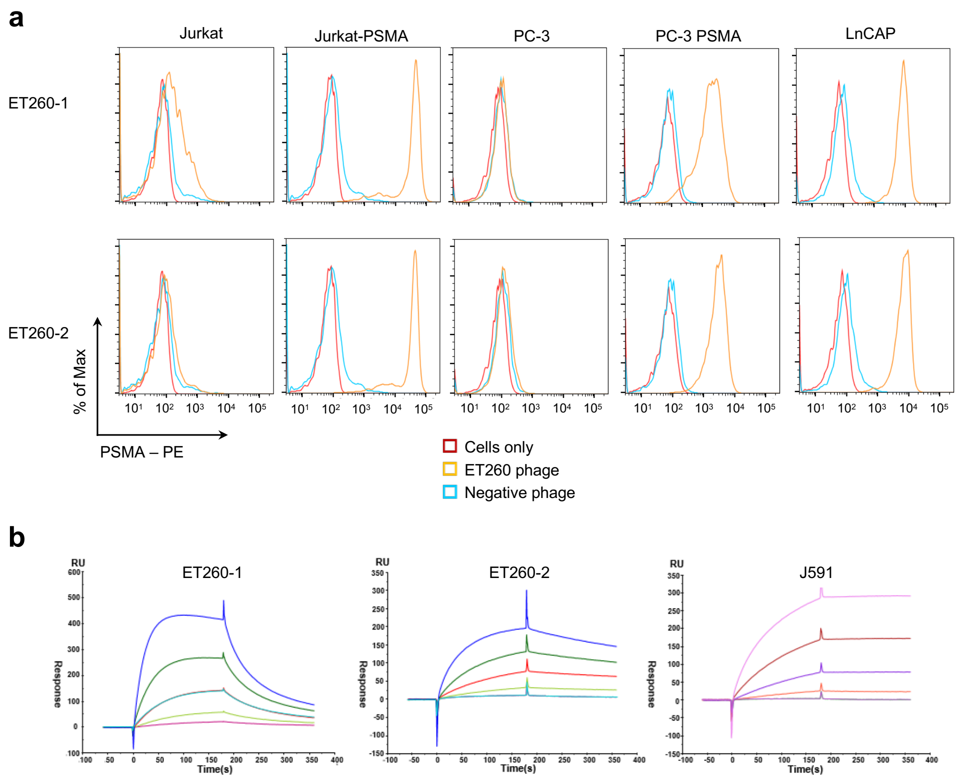
a
Jurkat
Jurkat-PSMA
PC-3
PC-3 PSMA
LnCAP
ET260-1
ET260-2
% of Max
PSMA – PE
Cells only
ET260 phage
Negative phage
b
ET260-1
ET260-2
J591
RU
Response
Time(s)

Figure S2

**a**

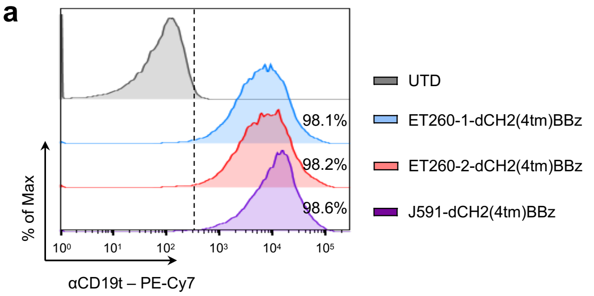

98.1%
98.2%
98.6%
UTD
ET260-1-dCH2(4tm)BBz
ET260-2-dCH2(4tm)BBz
J591-dCH2(4tm)BBz
% of Max
$10^0$ $10^1$ $10^2$ $10^3$ $10^4$ $10^5$
αCD19t – PE-Cy7


**b**

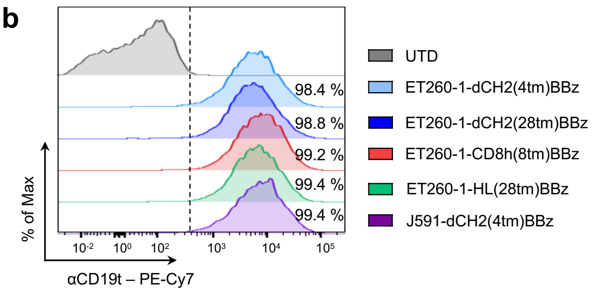

98.4 %
98.8 %
99.2 %
99.4 %
99.4 %
UTD
ET260-1-dCH2(4tm)BBz
ET260-1-dCH2(28tm)BBz
ET260-1-CD8h(8tm)BBz
ET260-1-HL(28tm)BBz
J591-dCH2(4tm)BBz
% of Max
$10^{-2}$ $10^0$ $10^2$ $10^3$ $10^4$ $10^5$
αCD19t – PE-Cy7

Figure S3

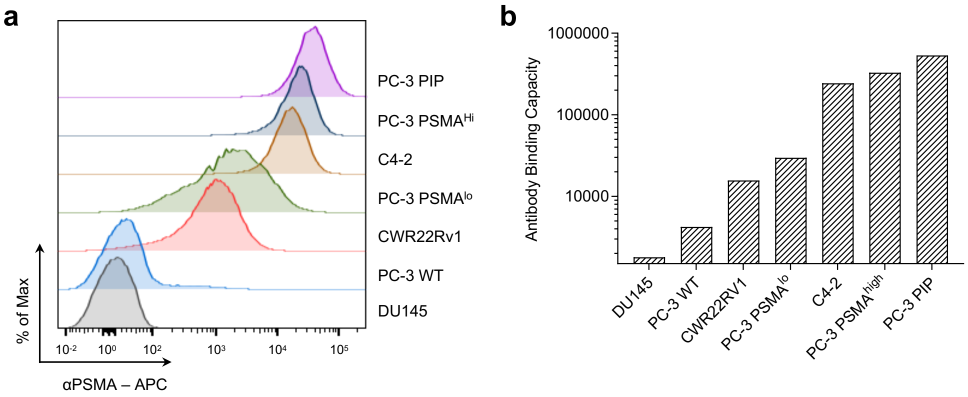
a
PC-3 PIP
PC-3 PSMA^Hi
C4-2
PC-3 PSMA^lo
CWR22Rv1
PC-3 WT
DU145
% of Max
αPSMA – APC
10^-2
10^0
10^2
10^3
10^4
10^5
b
Antibody Binding Capacity
1000000
100000
10000
DU145
PC-3 WT
CWR22RV1
PC-3 PSMA^lo
C4-2
PC-3 PSMA^high
PC-3 PIP

### Figure S4

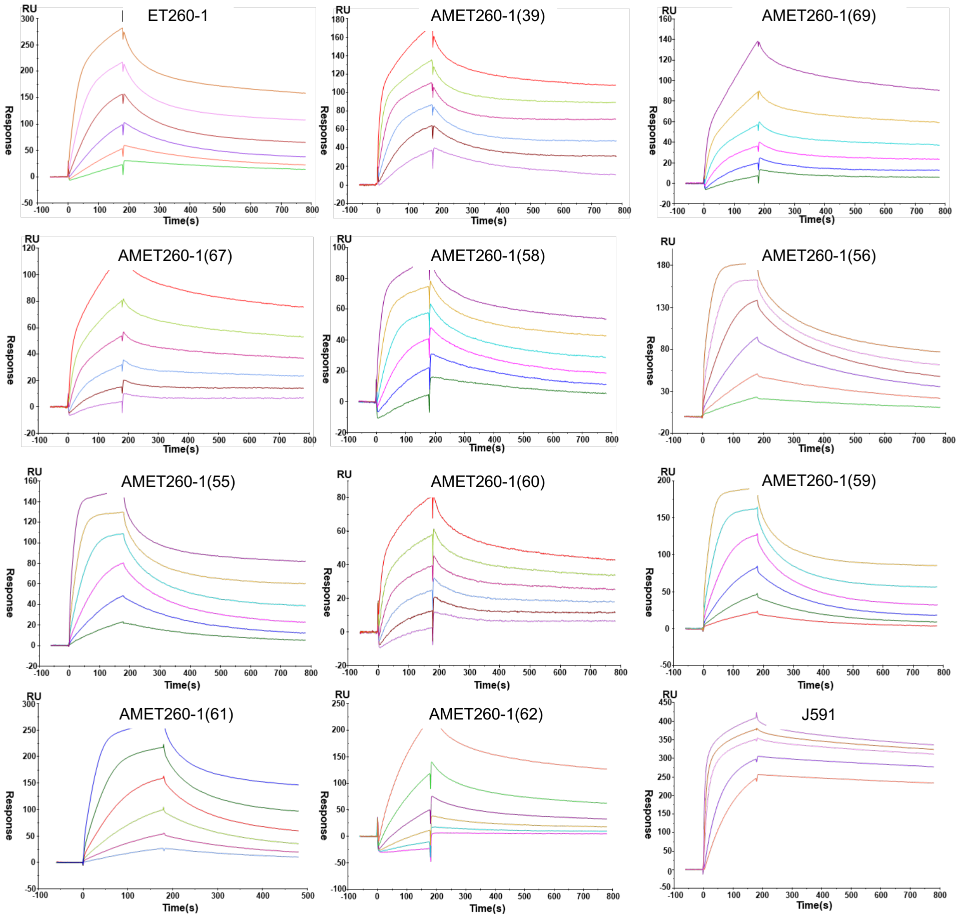
ET260-1
AMET260-1(39)
AMET260-1(69)
AMET260-1(67)
AMET260-1(58)
AMET260-1(56)
AMET260-1(55)
AMET260-1(60)
AMET260-1(59)
AMET260-1(61)
AMET260-1(62)
J591
RU
Response
Time(s)

Figure S5

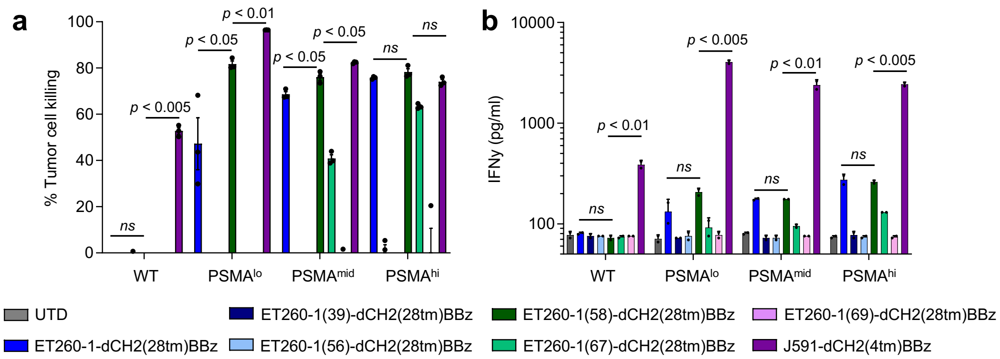
a
b
% Tumor cell killing
IFNγ (pg/ml)
ns
p < 0.005
p < 0.05
p < 0.01
p < 0.05
p < 0.05
ns
ns
ns
p < 0.01
ns
p < 0.005
ns
p < 0.01
ns
p < 0.005
WT
PSMAlo
PSMAmid
PSMAhi
UTD
ET260-1-dCH2(28tm)BBz
ET260-1(39)-dCH2(28tm)BBz
ET260-1(56)-dCH2(28tm)BBz
ET260-1(58)-dCH2(28tm)BBz
ET260-1(67)-dCH2(28tm)BBz
ET260-1(69)-dCH2(28tm)BBz
J591-dCH2(4tm)BBz

Figure S6

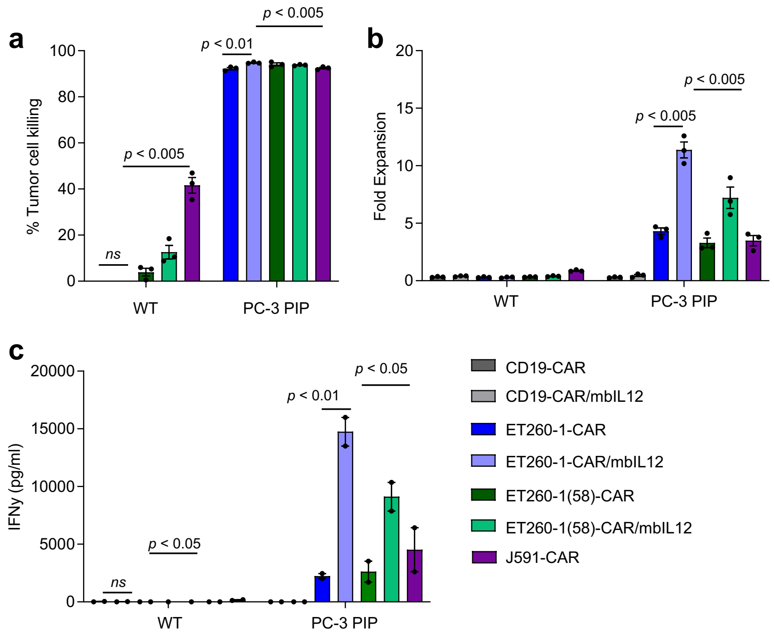

a
% Tumor cell killing
100
80
60
40
20
0
ns
p < 0.005
p < 0.01
p < 0.005
WT
PC-3 PIP
b
Fold Expansion
20
15
10
5
0
p < 0.005
p < 0.005
WT
PC-3 PIP
c
IFNγ (pg/ml)
20000
15000
10000
5000
0
ns
p < 0.05
p < 0.01
p < 0.05
WT
PC-3 PIP
CD19-CAR
CD19-CAR/mbIL12
ET260-1-CAR
ET260-1-CAR/mbIL12
ET260-1(58)-CAR
ET260-1(58)-CAR/mbIL12
J591-CAR

### Figure S7

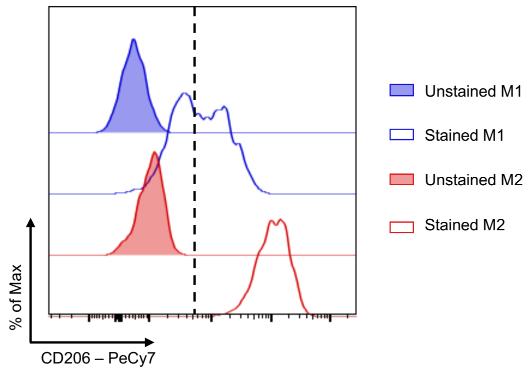

Unstained M1
Stained M1
Unstained M2
Stained M2
% of Max
CD206 – PeCy7

Figure S8

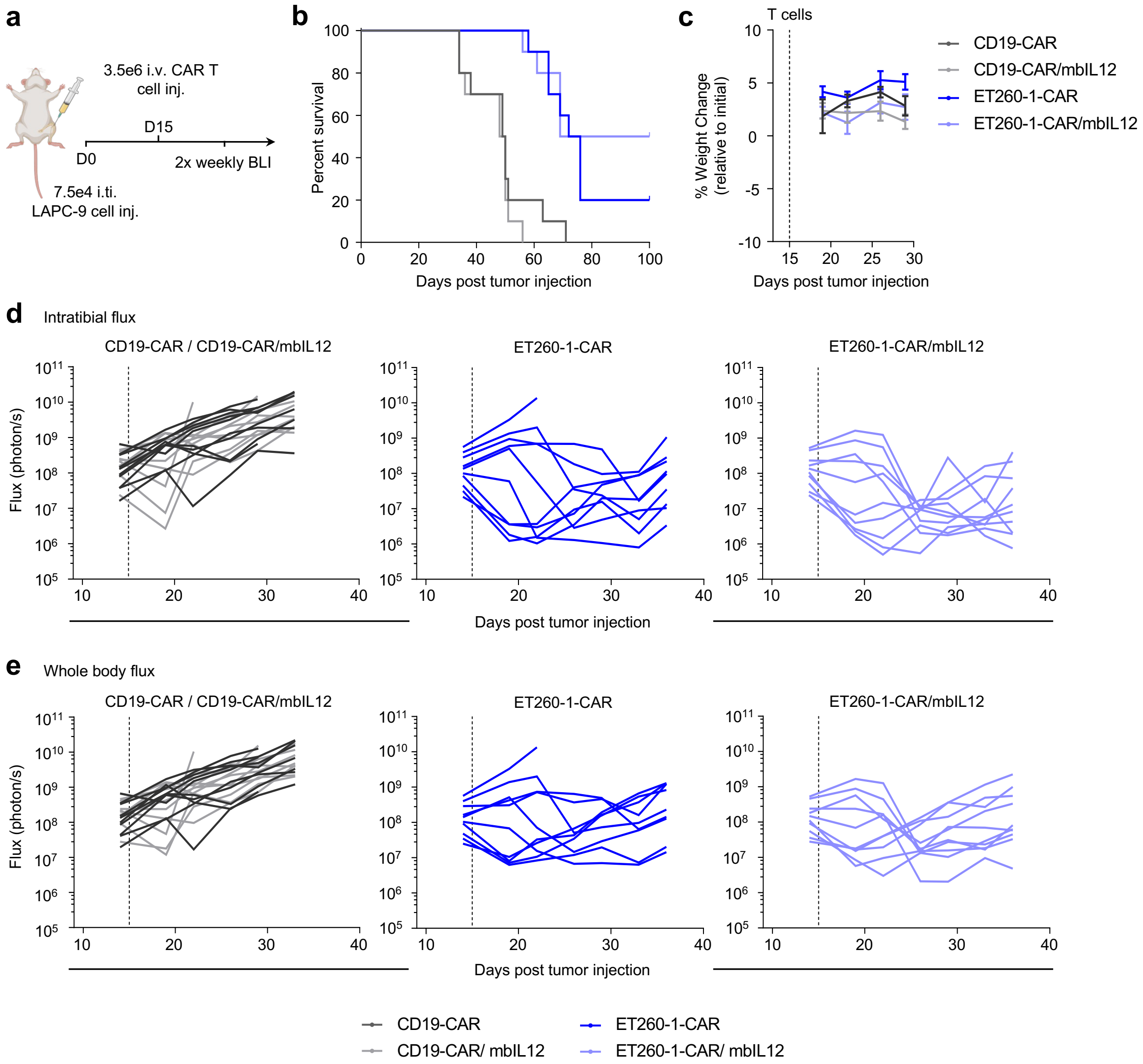

a
3.5e6 i.v. CAR T cell inj.
D15
D0
2x weekly BLI
7.5e4 i.ti. LAPC-9 cell inj.
b
Percent survival
Days post tumor injection
c
T cells
% Weight Change (relative to initial)
Days post tumor injection
CD19-CAR
CD19-CAR/mbIL12
ET260-1-CAR
ET260-1-CAR/mbIL12
d
Intratibial flux
CD19-CAR / CD19-CAR/mbIL12
ET260-1-CAR
ET260-1-CAR/mbIL12
Flux (photon/s)
Days post tumor injection
e
Whole body flux
CD19-CAR / CD19-CAR/mbIL12
ET260-1-CAR
ET260-1-CAR/mbIL12
Flux (photon/s)
Days post tumor injection
CD19-CAR
CD19-CAR/ mbIL12
ET260-1-CAR
ET260-1-CAR/ mbIL12

Table S1

| | k on (1/s µM) | k off (1/s) | KD (nM) |
|---|---|---|---|
| ET260-1 | 1.94 | 0.020 | 10.50 |
| ET260-2 | 0.64 | 0.002 | 2.37 |
| J591 | 0.33 | 0.00000045 | 0.0014 |

Table S2

| Sample Name | Antigen Binding Capacity |
|---|---|
| DU145 | 1,800 |
| PC-3 WT | 4,244 |
| CWR22RV1 | 15,725 |
| PC-3 $PSMA^{lo}$ | 29,722 |
| C4-2 | 244,072 |
| PC-3 $PSMA^{hi}$ | 329,100 |
| PC-3 PIP | 534,838 |

Table S3

| | **on-rate** ka1 (1/Ms) | **off-rate** kd1 (1/s) | KD1 (M) |
|---|---|---|---|
| ET260-1 | 1.97E+05 | 2.20E-03 | 1.120E-8 |
| AMET260-1-39 | 6.76E+05 | 5.42E-03 | 8.01E-09 |
| AMET260-1-69 | 1.37E+04 | 1.26E-04 | 9.19E-09 |
| AMET260-1-67 | 2.38E+04 | 4.99E-05 | 2.10E-09 |
| AMET260-1-58 | 3.61E+04 | 5.75E-05 | 1.60E-09 |
| AMET260-1-56 | 1.97E+05 | 4.01E-04 | 2.03E-09 |
| AMET260-1-55 | 3.57E+04 | 4.11E-04 | 1.15E-08 |
| AMET260-1-60 | 2.42E+05 | 4.93E-04 | 2.04E-08 |
| AMET260-1-59 | 3.66E+05 | 8.28E-04 | 2.27E-08 |
| AMET260-1-61 | 4.16E+05 | 1.92E-04 | 4.62E-08 |
| AMET260-1-62 | 1.43E+05 | 1.92E-03 | 4.62E-08 |
| J591 | 5.84E+05 | 2.04E-4 | 3.50E-10 |
